# Recounting the FANTOM Cage Associated Transcriptome

**DOI:** 10.1101/659490

**Authors:** Eddie-Luidy Imada, Diego Fernando Sanchez, Leonardo Collado-Torres, Christopher Wilks, Tejasvi Matam, Wikum Dinalankara, Aleksey Stupnikov, Francisco Lobo-Pereira, Chi-Wai Yip, Kayoko Yasuzawa, Naoto Kondo, Masayoshi Itoh, Harukazu Suzuki, Takeya Kasukawa, Chung-Chau Hon, Michiel JL de Hoon, Jay W Shin, Piero Carninci, FANTOM consortium, Andrew E Jaffe, Jeffrey T Leek, Alexander Favorov, Gloria R Franco, Ben Langmead, Luigi Marchionni

## Abstract

Long non-coding RNAs (lncRNAs) have emerged as key coordinators of biological and cellular processes. Characterizing lncRNA expression across cells and tissues is key to understanding their role in determining phenotypes including human diseases. We present here `FC-R2`, a comprehensive expression atlas across a broadly-defined human transcriptome, inclusive of over 109,000 coding and non-coding genes, as described in the FANTOM CAGE-Associated Transcriptome (FANTOM-CAT) study. This atlas greatly extends the gene annotation used in the original *recount2* resource. We demonstrate the utility of the `FC-R2` atlas by reproducing key findings from published large studies and by generating new results across normal and diseased human samples. In particular, we (a) identify tissue specific transcription profiles for distinct classes of coding and non-coding genes, (b) perform differential expression analyses across thirteen cancer types, providing new insights linking promoter and enhancer lncRNAs expression to tumor pathogenesis, and (c) confirm the prognostic value of several enhancers in cancer. Comprised of over 70,000 samples, the `FC-R2` atlas will empower other researchers to investigate functions and biological roles of both known coding genes and novel lncRNAs. Most importantly, access to the `FC-R2` atlas is available from https://jhubiostatistics.shinyapps.io/recount/, the *recount* Bioconductor package, and http://marchionnilab.org/fcr2.html.

## Introduction

Long non-coding RNAs (lncRNAs) are commonly defined as transcripts devoid of open reading frames (ORFs) longer than 200 nucleotides, which are often polyadenylated. This definition is not based on their function, since lncRNAs are involved in distinct molecular processes and biological contexts not yet fully characterized^1^. Over the past few years, the importance of lncRNAs has clearly emerged, leading to an increasing focus on decoding the consequences of their modulation, studying their involvement in the regulation of key biological mechanisms during development, normal tissue and cellular homeostasis, and in disease^1–3^.

Given the emerging and previously underestimated importance of non-coding RNAs, the FANTOM consortium has initiated the systematic characterization of their biological function. Through the use of Cap Analysis of Gene Expression sequencing (CAGE-seq), combined with RNA-seq data from the public domain, the FANTOM consortium released a comprehensive atlas of the human transcriptome, encompassing more accurate transcriptional start sites (TSS) for coding and non-coding genes, including numerous novel long non-coding genes: the FANTOM CAGE Associated Transcriptome (*FANTOM-CAT*)^4^. We hypothesized that these lncRNAs can be measured in many RNA-seq datasets from the public domain and that they have been so far missed by the lack of a comprehensive gene annotation.

Although the systematic analysis of lncRNAs function is being addressed by the FANTOM consortium in loss of function studies, increasing the detection rate of these transcripts combining different studies is difficult because the heterogeneity of analytic methods employed. Current resources that apply uniform analytic methods to create expression summaries from public data do exist but can miss several lncRNAs because their dependency on a pre-existing gene annotation for creating the genes expression summaries^5, 6^. We recently created *recount2*^7^, a collection of uniformly-processed human RNA-seq data, wherein we summarized 4.4 trillion reads from over 70,000 human samples from the Sequence Reads Archive (SRA), The Cancer Genome Atlas (TCGA)^8^, and the Genotype-Tissue Expression (GTEx)^9^ projects^7^. Importantly, *recount2* provides annotation-agnostic coverage files that allow re-quantification using a new annotation without having to re-process the RNA-seq data.

Given the unique opportunity to access lastest results to the most comprehensive human transcriptome (the *FANTOM-CAT* project) and the *recount2* gene agnostic summaries, we addressed the previous described challenges building a comprehensive atlas of coding and non-coding gene expression across the human genome: the *FANTOM-CAT/recount2* expression atlas (`FC-R2` hereafter). Our resource contains expression profiles for 109,873 putative genes across over 70,000 samples, enabling an unparalleled resource for the analysis of the human coding and non-coding transcriptome.

## Results

### Building the *FANTOM-CAT/recount2* resource

The *recount2* resource includes a coverage track, in the form of a BigWig file, for each processed sample. We built the `FC-R2` expression atlas by extracting expression levels from *recount2* coverage tracks in regions that overlapped unambiguous exon coordinates for the permissive set of *FANTOM-CAT* transcripts, according to the pipeline shown in Figure 1. Since *recount2*’s coverage tracks does not distinguish from between genomic strands, we removed ambiguous segments that presented overlapping exon annotations from both strands (see Methods section and Supp. Methods). After such disambiguation procedure, the remaining 1,066,515 exonic segments mapped back to 109,869 genes in *FANTOM-CAT* (out of the 124,047 starting ones included in the permissive set^4^). Overall, the `FC-R2` expression atlas encompasses 2,041 studies with 71,045 RNA-seq samples, providing expression information for 22,116 coding genes and 87,763 non-coding genes, such as enhancers, promoters, and others lncRNAs.

**Figure 1.**
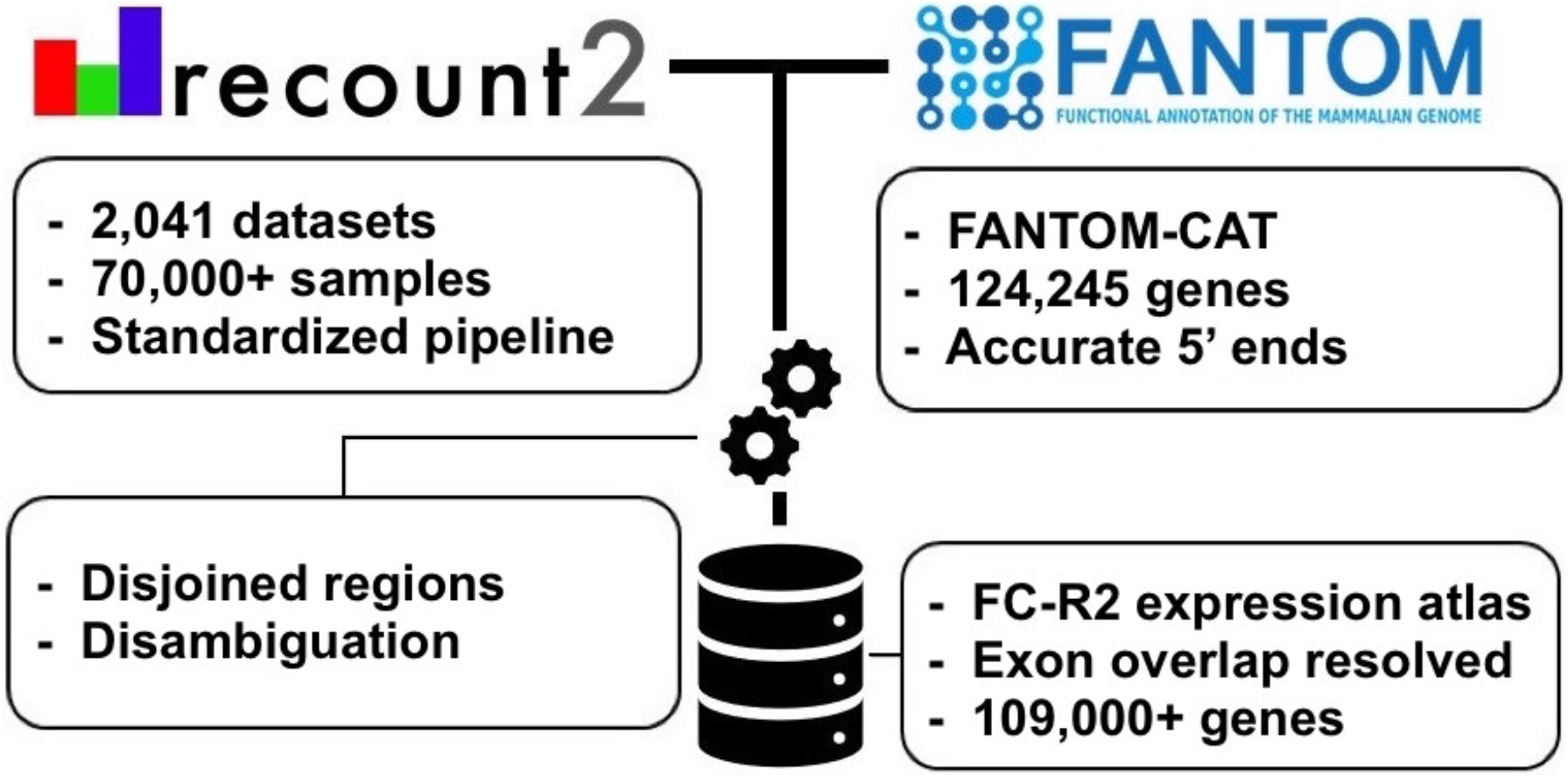
Overview of the *FANTOM-CAT/recount2* resource development. `FC-R2` leverages two public resources, the *FANTOM-CAT* gene models and *recount2*. `FC-R2` provides expression information for 109,873 genes, both coding (22,110) and non-coding (87,693). This latter group encompasses enhancers, promoters, and others lncRNAs.

### Validating the *FANTOM-CAT/recount2* resource

We first assessed how gene expression estimates in `FC-R2` compared to previous gene expression estimates from other projects. Specifically, we considered data from the GTEx consortium (v6), spanning 9,662 samples from 551 individuals and 54 tissues types^9^. First, we computed the correlation for the GTEx data between gene expression based on the `FC-R2` atlas and on GENCODE (v25) gene model in *recount2*, which has been already shown to be consistent with gene expression estimates from the GTEx project^7^, observing a median correlation ≥ 0.986 for the 32,922 genes in common. This result supports the notion that our pre-processing steps to disambiguate overlapping exon regions between strands did not significantly alter gene expression quantification.

Next, we assessed whether gene expression specificity, as measured in `FC-R2`, was maintained across tissue types. To this end, we selected and compared gene expression for known tissue-specific expression patterns, such as Keratin 1 (*KRT1*), Estrogen Receptor 1 (*ESR1*), and Neuronal Differentiation 1 (*NEUROD1*) (Figure 2). Overall, all analyzed tissue specific markers presented nearly identical expression profiles across GTEx tissue types between the alternative gene models considered (see Figure 2 and S1), confirming the consistency between gene expression quantification in `FC-R2` and those based on GENCODE.

**Figure 2.**
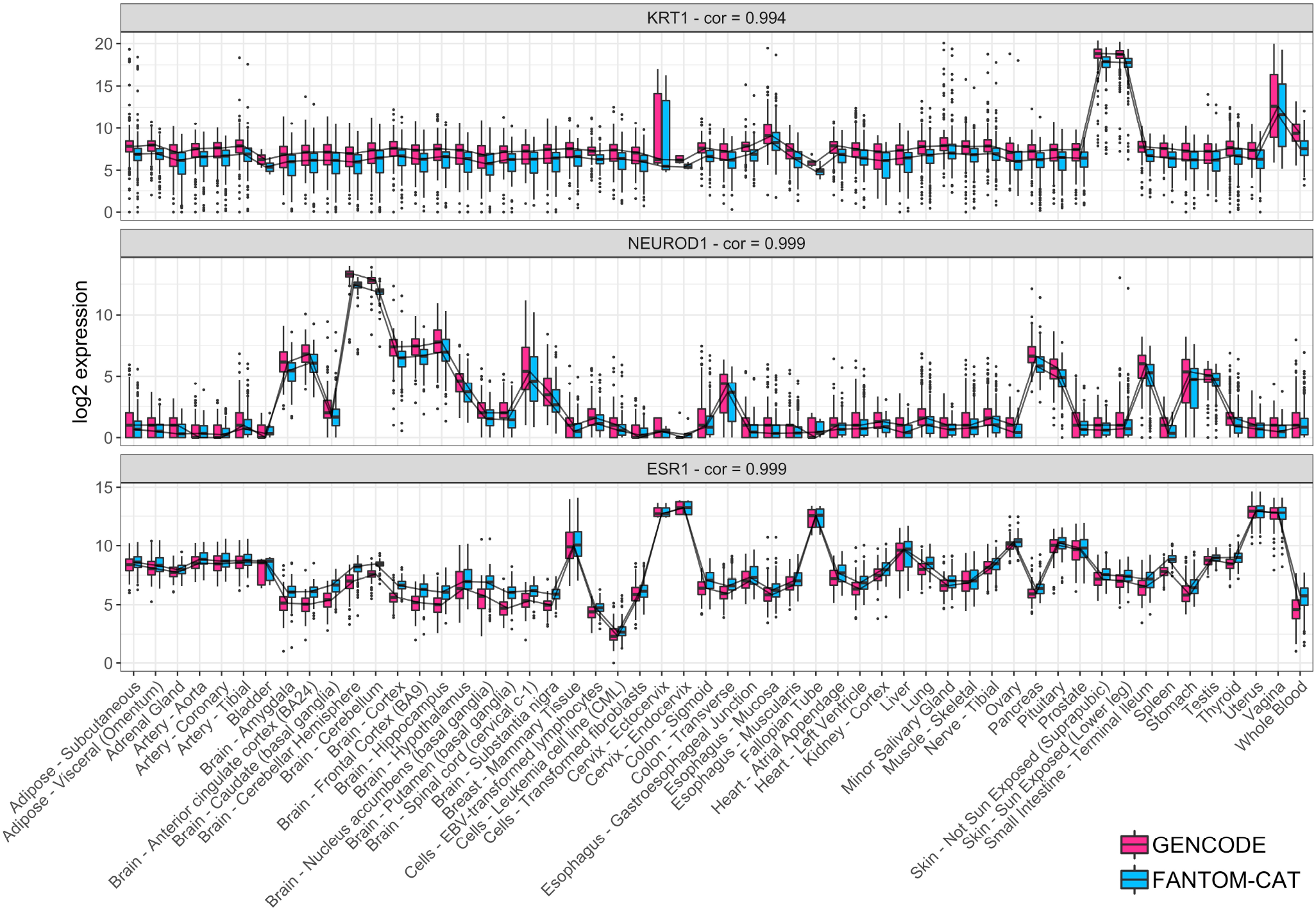
Tissue specific expression in GTEx. Log2 expression for three tissue specific genes (*KRT1*, *NEUROD1*, and *ESR1*) in GTEx data stratify by tissue type using `FC-R2` and GENCODE based quantification. Expression profiles are highly correlated and expressed consistently in the expected tissue types (*e.g.*, *KRT1* is most expressed in skin, *NEUROD1* in brain, and *ESR1* in estrogen sensitive tissue types like uterus, Fallopian tubes, and breast). Correlations are shown on top for each tissue marker. Center lines, upper/lower quartiles and Whiskers represents the median, 25/75 quartiles and 1.5 interquartile range, recpectively.

### Tissue-specific expression of lncRNAs

It has been shown that, although expressed at a lower level, enhancers and promoters are not ubiquitously expressed and are more specific for different cell types than coding genes^4^. In order to verify this finding, we used GTEx data to assess expression levels and specificity profiles across samples from each of the 54 analyzed tissue types, stratified into four distinct gene categories: coding mRNA, intergenic promoter lncRNA (ip-lncRNA), divergent promoter lncRNA (dp-lncRNA), and enhancers lncRNA (e-lncRNA). Overall, we were able to confirm that these RNA classes are expressed at different levels, and that they display distinct specificity patterns across tissues, as shown for primary cell types by Hon et al.^4^, albeit with more variability likely due to the increased cellular complexity present in tissues. Specifically, coding mRNAs were expressed at higher levels than lncRNAs (log2 median expression of 6.6 for coding mRNAs, and of 4.1, 3.8 and 3.1, for ip-lncRNA, dp-lncRNA, and e-lncRNA, respectively). In contrast, the expression of enhancers and intergenic promoters was more tissue-specific (median = 0.41 and 0.30) than what observed for divergent promoters and coding mRNAs (median = 0.13 and 0.09) (Figure 3A). Finally, when analyzing the percentage of genes expressed across tissues by category, we observed that coding genes are, in general, ubiquitous, while lncRNAs are more specific, with enhancers showing the lowest percentages of expressed genes (mean ranging from 88.42% to 41.98%, see Figure 3B), in agreement with the notion that enhancer transcription is tissue specific^10^.

**Figure 3.**
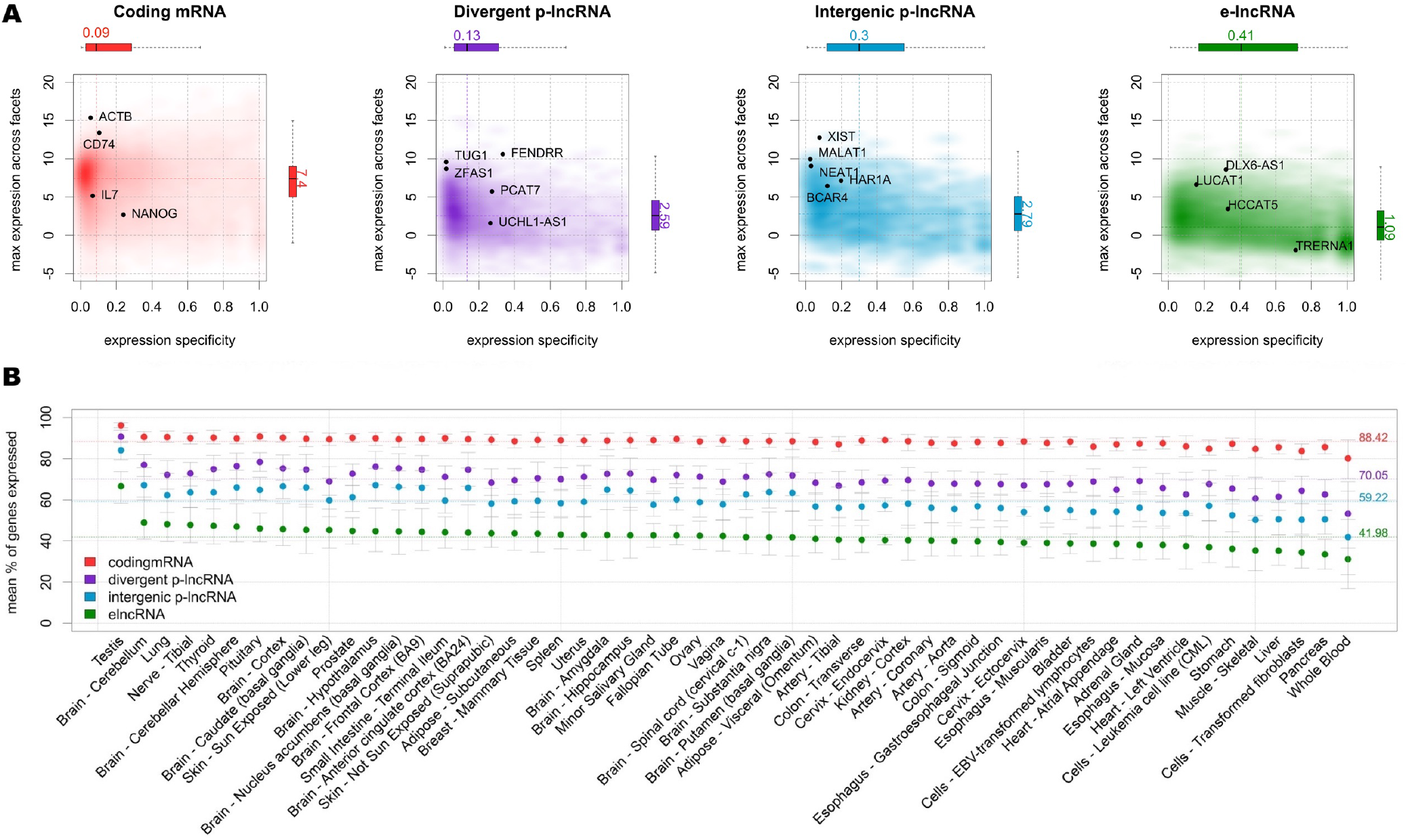
Expression profiles across GTEx tissues. **A)** Expression level and tissue specificity across four distinct RNA categories. The Y-axis shows log2 expression levels representing each gene using its maximum expression in GTEx tissues expressed as transcripts per million (TMP). The X-axis shows expression specificity based on entropy computed from median expression of each gene across the GTEx tissue types. Individual genes are highlighted in the figure panels. **B)** Percentage of genes expressed for each RNA category stratified by GTEx tissue facets. The dots represent the mean among samples within a facet and the error bars represent 99.99% confidence intervals. Dashed lines represent the means among all samples.

### Differential expression analysis of coding and non-coding genes in cancer

We analyzed coding and non-coding genes expression in cancer using TCGA data. To this end, we compared cancer to normal samples separately for 13 tumor types, using `FC-R2` re-quantified data. We further identified the differentially expressed genes (DEG) in common across the distinct cancer types (see Figure 4). Overall, the number of DEG varied across cancer types and by gene class, with a higher number of significant coding than non-coding genes (FDR < 0.01, see table 1). Importantly, a substantial fraction of these genes was exclusively annotated in the *FANTOM-CAT*, suggesting that relying on other gene models would result in missing many potential important genes (see Table 1). We then analyzed the consensus among cancer types. A total of 41 coding mRNAs were differentially expressed across all the 13 tumor types after global correction for multiple testing (FDR < 10^−6^, see Supplementary table S1). For lncRNAs, a total of 28 divergent promoters, four intergenic promoters, and three enhancers were consistently up- or down-regulated across all the 13 tumor types after global correction for multiple testing (FDR < 0.1, see Supplementary tables S2, S3, S4, respectively).

**Figure 4.**
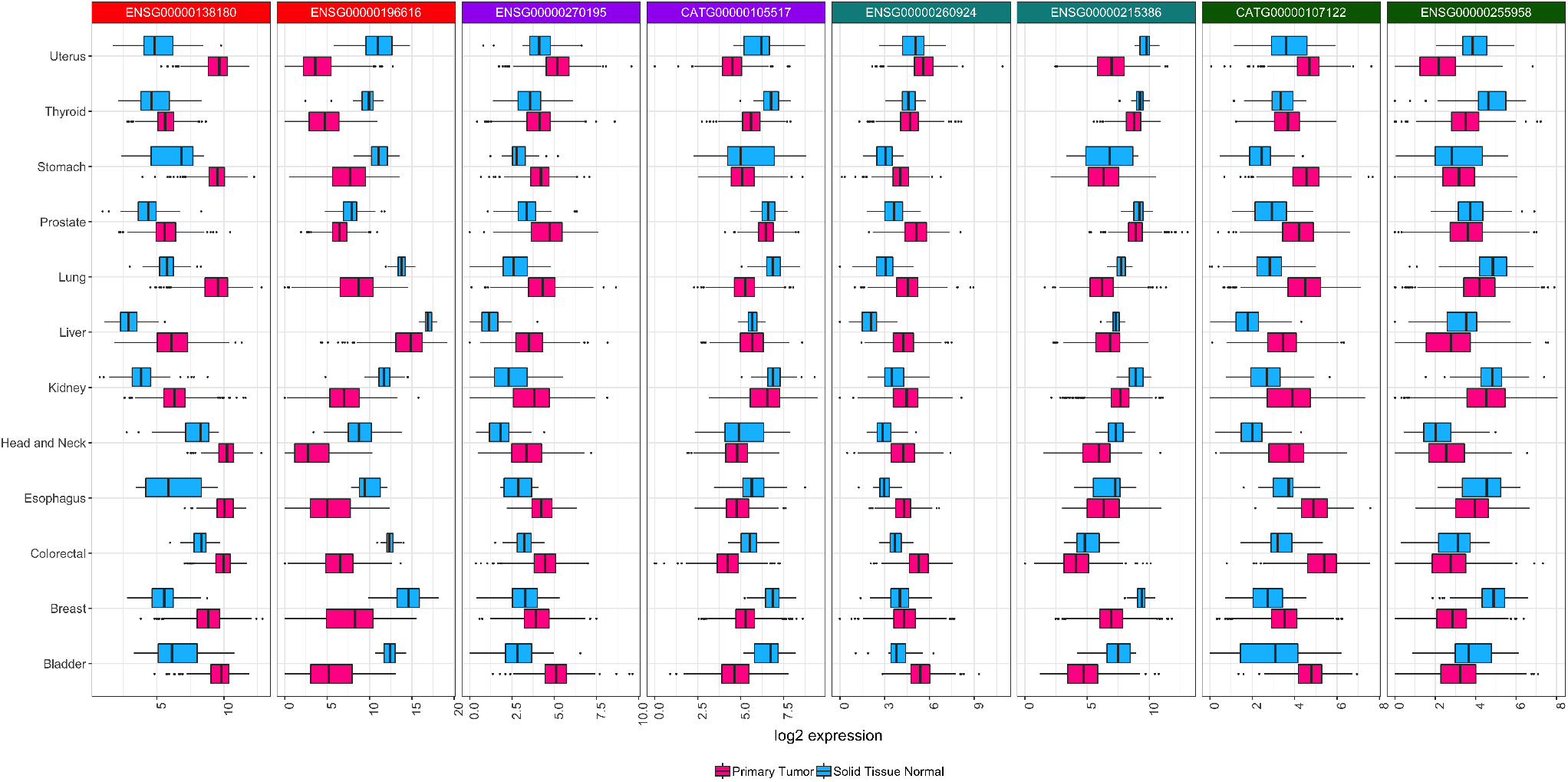
Differential expression for selected transcripts from distinct RNA classes across tumor types. Boxplots for selected differential expressed genes between tumor and normal samples across all 13 tumor types analyzed. For each tissue of origin, the most up-regulated (on the left) and down-regulated (on the right) gene for each RNA class is shown. Center lines, upper/lower hinges, and the whiskers respectively represent the median, the upper and lower quartiles, and 1.5 extensions of the interquartile range. Color coding on top of the figure indicates the RNA classes (red for mRNA, purple for dp-lncRNA, cyan ip-lncRNA, and green for e-lncRNA. These genes were select after global multiple testing correction across all 13 tumor types (see Supplementary Tables S1, S2, S3, and S4)

**Table 1.** Differentially expressed genes in cancer. The table below summarizes the number of significant DEG (*FDR* ≤ 0.01) between tumor and normal samples across the 13 cancer types analyzed for each gene class considered (coding mRNA, ip-lncRNA, dp-lncRNA, and e-lncRNA). Counts are reported separately for DEG up- and down-regulated in cancer, and values in parenthesis represents the number of genes exclusively annotated in the *FANTOM-CAT* gene model. Mean and standard deviation across cancer types is shown at the bottom.

| Cancer type | Total | dp-lncRNA |  | e-lncRNA |  | ip-lncRNA |  | mRNA |  |
| --- | --- | --- | --- | --- | --- | --- | --- | --- | --- |
|  |  | Up | Down | Up | Down | Up | Down | Up | Down |
| Bile | 7010 | 200 (60) | 313 (90) | 186 (89) | 203 (99) | 47 (12) | 84 (17) | 2658 (106) | 3319 (97) |
| Bladder | 7680 | 344 (125) | 319 (87) | 140 (68) | 149 (67) | 65 (19) | 82 (7) | 3112 (201) | 3469 (61) |
| Breast | 15290 | 753 (291) | 721 (202) | 656 (377) | 583 (305) | 207 (50) | 178 (32) | 6109 (296) | 6083 (244) |
| Colorectal | 13685 | 490 (164) | 592 (168) | 381 (203) | 400 (196) | 130 (32) | 160 (28) | 5538 (371) | 5994 (132) |
| Esophagus | 4883 | 87 (21) | 193 (50) | 90 (38) | 184 (103) | 40 (11) | 48 (2) | 1921 (83) | 2320 (77) |
| Head and Neck | 10517 | 442 (138) | 401 (96) | 267 (139) | 251 (112) | 100 (23) | 109 (18) | 4329 (256) | 4618 (53) |
| Kidney | 15697 | 734 (238) | 820 (281) | 535 (299) | 486 (209) | 203 (45) | 200 (48) | 6349 (525) | 6370 (114) |
| Liver | 10554 | 346 (94) | 395 (106) | 230 (102) | 248 (123) | 90 (16) | 112 (19) | 4164 (174) | 4969 (95) |
| Lung | 17143 | 864 (338) | 835 (304) | 893 (512) | 729 (396) | 242 (76) | 213 (39) | 7523 (532) | 5844 (212) |
| Prostate | 13183 | 686 (287) | 654 (218) | 418 (254) | 452 (214) | 175 (55) | 167 (30) | 5153 (489) | 5478 (128) |
| Stomach | 11309 | 528 (213) | 518 (164) | 462 (291) | 436 (240) | 144 (51) | 129 (22) | 4509 (558) | 4583 (89) |
| Thyroid | 14264 | 752 (284) | 804 (318) | 527 (295) | 594 (332) | 161 (39) | 174 (47) | 5403 (189) | 5849 (308) |
| Uterus | 12906 | 641 (285) | 713 (235) | 454 (263) | 612 (341) | 210 (79) | 225 (54) | 5135 (335) | 4916 (181) |
| Mean | 11855 | 528 (195) | 560 (178) | 403 (225) | 410 (211) | 140 (39) | 145 (28) | 4762 (317) | 4909 (138) |
| St. Dev | 3650 | 237 (102) | 218 (89) | 225 (137) | 189 (107) | 67 (23) | 55 (16) | 1557 (167) | 1234 (77) |

Next, we reviewed the literature to assess functional correlates for such consensus genes. Most of the up-regulated coding genes (Supplementary Table S1) participate in cell cycle regulation, cell division, DNA replication and repair, chromosome segregation, and mitotic spindle checkpoints. Most of the consensus down-regulated mRNAs (Supplementary Table S1) are associated with metabolism and oxidative stress, transcriptional regulation, cell migration and adhesion, and with modulation of DNA damage repair and apoptosis.

Three down-regulated dp-lncRNA genes, *RP11-276H19*, *RPL34-AS1*, and *RAP2C-AS1*, were reported to be implicated in cancer (Supplementary Table S2). The first one controls epithelial-mesenchymal transition, the second is associated with tumor size increase, while the third is associated with urothelial cancer after kidney cancer transplantation^11–13^. Among the up-regulated dp-lncRNAs (Supplementary Table S2), *SNHG1* has been implicated in cellular proliferation, migration and invasion of different cancer types, and to be strongly up-regulated in osteosarcoma, non-small lung cancer, and gastric cancer^14, 15^.

Among the ip-lncRNAs ubiquitously down-regulated (see Supplementary Table S3), *LINC00478* has been identified in many different tumors types including leukemia, breast, vulvar, prostate, and bladder cancer^16–20^. For instance, in vulvar squamous cell carcinoma, *LINC00478* and *MIR31HG* expressions are correlated and associated with tumor differentiation^17^. Similarly, *LINC00478* down-regulation in ER positive breast cancer is associated with progression, recurrence, and metastasis^18^. In contrast, increased expression of *SNHG17* (an ip-lncRNA, see Supplementary Table S3), was associated with short term survival in breast cancer, and with tumor size, stage, and lymph node metastasis in colorectal cancer^21, 22^. In addition, *AC004463*, another ip-lncRNA (Supplementary Table S3), was found to be up-regulated in liver cancer and metastatic prostate cancer^23^. Despite we did not identify any cancer association for common e-lncRNAs, one among those we identified, *RP5-965F6*, has been previously reported to be up-regulated in late-onset Alzheimer’s disease^24^. Furthermore, the enhancers lncRNA class also yielded the lowest number of genes in common among all cancer types, reinforcing the concept that enhancers are expressed in a tissue specific manner (See Figure 3A and Supplementary Table S4).

Finally, we focused more in depth on prostate cancer (PCa) as a prototypical example, and we were able to confirm previous findings for both coding and non-coding genes (see Supplementary Figure S2). For coding genes, we confirmed differential expression for known markers of PCa progression and mortality, like *ERG*, *FOXA1*, *RNASEL*, *ARVCF*, and *SLC43A1*^25, 26^. Similarly, we also confirmed differential expression for non-coding genes, like *PCA3*, the first clinically approved lncRNA marker for PCa^27, 28^, *PCAT1*, a prostate-specific lncRNA involved in disease progression^29^, *MALAT1*, which is associated with PCa poor prognosis^30^, *CDKN2B-AS1*, an anti-sense lncRNA up-regulated in PCa that inhibits tumor suppressor genes activity^31, 32^, and the *MIR135* host gene, which is associated with castration-resistant PCa^33^.

### Enhancer expression levels hold prognostic value

The number of lncRNAs involved in cancer development and progression is rapidly increasing, we therefore analyzed the prognostic value of the lncRNAs we identified in our gene expression differential analysis in TCGA, as well as those previously reported in other studies. To this end, Chen and collaborators have recently surveyed enhancers expression in nearly 9,000 patients from TCGA^34^, using genomic coordinates from the FANTOM5 project^35^, identifying 4,803 enhancers with prognostic potential in one or more TCGA tumor types. We therefore leveraged the `FC-R2` atlas to identify prognostic coding and non-coding genes using Univariate Cox proportional hazard models, comparing our results for e-lncRNAs with those reported by Chen and colleagues.

When we considered e-lncRNA expression, we identified a total of 5,382 prognostic e-lncRNAs (FDR ≤ 0.05), and no single one was associated with survival across all cancer types. Overall, the number of significant prognostic e-lncRNAs varied across tumor types, ranging from 3 in head and neck cancer to 3,850 in kidney cancers (see Supplementary Table S6). Notably, two (out of three) e-lncRNAs from our differential gene expression consensus list across all tumor types were also prognostic. Higher expression of *CATG00000107122* gene was associated with worse prognosis in kidney cancer (Supplementary figure S4b) Overall, despite differences in annotation and quantification (see Supplementary Table S5), we were able to confirm prognostic value for 2,765 e-lncRNAs out of the 4,803 reported by Chen et al^34^, including *“enhancer 22”* (*ENSG00000272666*), which was highlighted as a promising prognostic marker for kidney cancer (Supplementary Figure S3).

Finally, we analyzed the prognostic value for dp-lncRNAs, ip-lncRNAs, and mRNAs (See Supplementary Tables S7, S8, and S9, respectively), and assessed the survival prognostic potential of our consensus genes across tumor types. Thirty-seven of the 41 coding mRNAs, 22 of the 28 differentially expressed dp-lncRNAs, and two out of the four DE ip-lncRNAs, respectively, were found to be prognostic (See Supplementary Tables S10, S11, S12, and S13). Kaplan-Meier survival curves for one selected DE gene on each RNA subtype evaluated here are shown in supplementary figure S4.

## Discussion

The importance of lncRNAs in cell biology and disease has clearly emerged in the past few years and different classes of lncRNAs have been shown to play crucial roles in cell regulation and homeostasis^36^. For instance, enhancers – a major category of gene regulatory elements, which has been shown to be expressed^35, 37^ – play a prominent role in oncogenic processes^38, 39^ and other human diseases^40, 41^. Despite their importance, however, there is a scarcity of large-scale datasets investigating enhancers and other lncRNA classes, in part due to the technical difficulty in applying high-throughput techniques such as ChIP-seq and Hi-C over large cohorts, and to the use of gene models that do not account for them in transcriptomics analyses. Furthermore, the large majority of the lncRNAs that are already known – and that have been shown to be associated with some phenotype – are still lacking functional annotation.

To address these needs, the FANTOM consortium has first constructed the *FANTOM-CAT* meta-transcriptome, a comprehensive atlas of coding and non-coding genes with robust support from CAGE-seq data^4^, then it has undertaken a large scale project to systematically target lncRNAs and characterize their function using a multi-pronged approach (Ramilowski J, et al., manuscript in review^42^). In a complementary effort, we have leveraged public domain gene expression data from *recount2*^7, 43^ to create a comprehensive gene expression compendium across human cells and tissues based on the *FANTOM-CAT* gene model, with the ultimate goal of facilitating lncRNAs annotation through association studies. To this end, the `FC-R2` atlas is already in use in the FANTOM6 project (http://fantom.gsc.riken.jp/6/) to successfully characterize lncRNA expression in human samples (Ramilowski J, et al., manuscript in review^42^).

In order to validate our resource, we have compared the gene expression summaries based on *FANTOM-CAT* gene models with previous, well-established gene expression quantifications, demonstrating virtually identical profiles across tissue types overall and for specific tissue markers. We have then confirmed that distinct classes of coding and non-coding genes differ in terms of overall expression levels and specificity patterns across cell types and tissues. Furthermore, with this approach, we were also able to identify mRNAs, promoters, enhancers, and other lncRNAs that are differentially expressed in cancer, both confirming previously reported findings, and identifying novel cancer genes exclusively annotated in the *FANTOM-CAT* gene model, which have been therefore missed in prior analyses with TCGA data. Finally, we also analyzed the prognostic value of the coding and non-coding genes we identified, and confirmed survival associations in TCGA for measurable enhancers.

Collectively, by confirming findings reported in previous studies, our results demonstrate that the `FC-R2` gene expression atlas is a reliable and powerful resource for exploring both the coding and non-coding transcriptome, providing compelling evidence and robust support to the notion that lncRNA gene classes, including enhancers and promoters, despite not being yet fully understood, portend significant biological functions. Our resource, therefore, constitutes a suitable and promising platform for future large scales studies in cancer and other human diseases, which in turn hold the potential to reveal important cues to the understanding of their biological, physiological, and pathological roles, potentially leading to improved diagnostic and therapeutic interventions.

Finally, all results, data, and code from the `FC-R2` atlas are available as a public tool. With uniformly processed expression data for over 70,000 samples and 109,873 genes ready to analyze, we want to encourage researchers to dive deeper into the study of ncRNAs, their interaction with coding and non-coding genes, and their influence on normal and disease tissues. We hope this new resource will help paving the way to develop new hypotheses that can be followed to unwind the biological role of the transcriptome as a whole.

## Methods

### Data and pre-processing

The FANTOM CAT gene catalog (permissive set) was obtained from the FANTOM consortium within the frame of the FANTOM6 project (Ramilowski J, et al., manuscript in review^42^). This catalog accounts for 124,245 genes supported by CAGE peaks and it includes those described by Hon et al.^4^. In order to remove ambiguity due to overlapping among exons from distinct genes, the BED files containing the coordinates for all genes and exons were processed with the *GenomicRanges* R/Bioconductor package^44^ to obtain exon coordinates disjoining. To avoid losing strand information from annotation we processed data using a two-step approach by first disjoining overlapping segments on the same strand and then across strands (Figure 5). The genomic ranges (disjoint exons segments) that mapped back to more than one gene were discarded. The expression values for these ranges were then quantified using *recount.bwtool*^45^ (code at https://github.com/LieberInstitute/marchionni_projects). The resulting expression quantification were processed to generate `RangedSummarizedExperiment` objects compatible with the *recount2* framework^7, 43^ (code available from https://github.com/eddieimada/fcr2). Thus, the `FC-R2` atlas provides expression information for coding and non-coding genes (including enhancers, divergent promoters, and intergenic lncRNAs) for 9,662 samples from the Genotype-Tissue Expression (GTEx) project, 11,350 samples from TCGA, and over 50,000 samples from the Sequence Read Archive (SRA).

**Figure 5.**
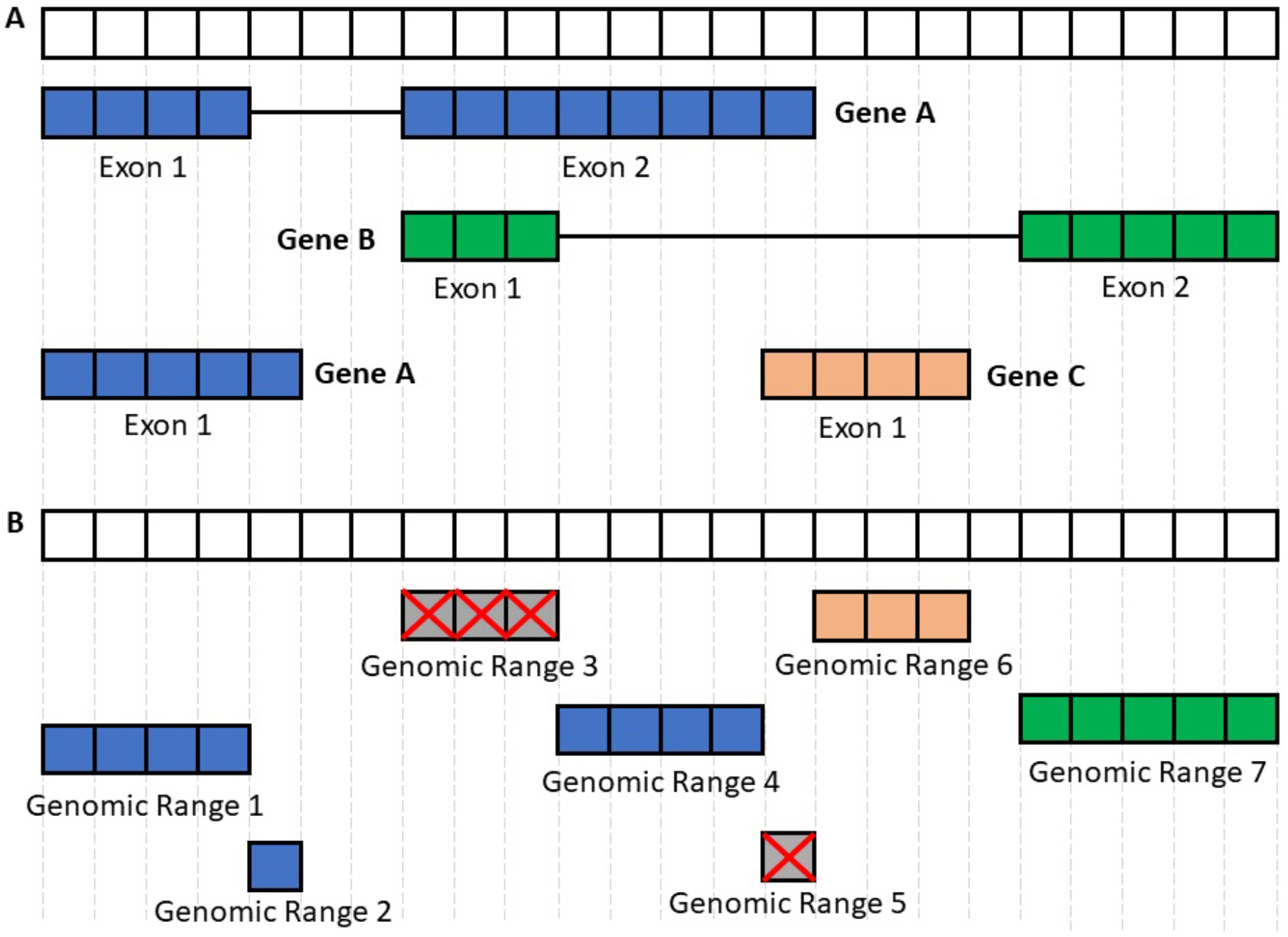
Processing the FANTOM-CAT genomic ranges. This figure summarizes the disjoining and exon disambiguation processes performed before extracting expression information from *recount2* using the *FANTOM-CAT* gene models. **A)** Representation of a genomic segment containing 3 distinct, hypothetical genes: gene A having two isoforms, and genes B and C with one isoform each. Each box can be interpreted as one nucleotide along the genome. Colors indicate the 3 different genes. **B)** Representation of disjoint exon ranges from example in Panel A. Each feature is reduced to a set of non-overlapping genomic ranges. The disjoint genomic ranges mapping back to two or more distinct genes are removed (crossed grey boxes). After removal of ambiguous ranges, the expression information for remaining ones is extracted from *recount2* and summarized at the gene level.

### Correlation with other studies

To test if the pre-processing steps used for `FC-R2` had a major impact on gene expression quantification, we compared our data to the published GTEx expression values obtained from *recount2* (version 2, https://jhubiostatistics.shinyapps.io/recount/). Specifically, we first compared the expression distribution of tissue specific genes across different tissue types and then computed the Pearson correlation for each gene in common across the original *recount2* gene expression estimates based on GENCODE and our version based on the FANTOM-CAT transcriptome.

### Expression specificity of tissue facets

We analyzed the expression level and specificity of each gene stratified by RNA class (i.e. mRNA, e-lncRNA, dp-lncRNA, ip-lncRNA) using the same approach described by Hon et al.^4^ (see Supplementary Methods). Briefly, overall expression levels for each gene were represented by the maximum transcript per million (TPM) values observed across all samples within each tissue type in GTEx. Gene specificity was based on the empirical entropy computed using the mean expression value across tissue types. The 99.99 percent confidence intervals for the expression of each category by tissue type were calculated based on TPM values. Genes with a TPM greater than 0.01 were considered expressed.

### Identification of differentially expressed genes

We analyzed differential gene expression in 13 cancer types, comparing primary tumor with normal samples using TCGA data from the `FC-R2` atlas. Gene expression summaries for each cancer type were split by RNA class (coding mRNA, intergenic promoter lncRNA, divergent promoter lncRNA, and enhancer lncRNA) and then analyzed independently. A generalized linear model approach coupled with empirical Bayes moderation of standard errors^46^ was used to identify differentially expressed genes between groups. The model was adjusted for the three most relevant coefficients for data heterogeneity as estimated by surrogate variable analysis (SVA)^47^. Correction for multiple testing was performed across RNA classes by merging the resulting p-values for each cancer type and applying the Benjamini-Hochberg method^48^.

### Prognostic analysis

To evaluate the prognostic potential of the genes in `FC-R2`, we performed univariate Cox proportional regression analysis separately for each RNA classes (22106 mRNAs, 17,404 e-lncRNAs, 6,204 dp-lncRNAs, and 1,948 ip-lncRNAs) across each of the 13 TCGA cancer types with available survival follow-up. Genes with FDR equal or less than 0.05, using Benjamini-Hochberg correction^48^ within the cancer type and RNA class, were deemed as significant prognostic factors. We further analyzed the prognostic value of the consensus genes we identified comparing tumors to normal samples by intersecting the corresponding gene lists with those obtained by Cox proportional regression. Finally, in order to compare the results from prognostic analyses, we obtained data on enhancers position and prognostic potential from Chen et al. original publication^34^ and performed a liftover to the hg38 genome assembly to match `FC-R2` coordinates.

## Data Availability

All data is available from http://marchionnilab.org/fcr2.html. Expression data can be directly accessed through https://jhubiostatistics.shinyapps.io/recount/ and the *recount* Bioconductor package (v1.9.5 or newer) at https://bioconductor.org/packages/recount as *RangedSummarizedExperiment* objects organized by The Sequence Read Archive (SRA) study ID. The data can be loaded using R-programming language and is ready to be analyzed using Bioconductor packages or the data can be exported to other formats for use in another environment.

## Code Availability

All code used in this manuscript is available for reproducibility and transparency at: https://github.com/eddieimada/fcr2 and https://github.com/LieberInstitute/marchionni_projects.

## Acknowledgements

This publication was made possible though support from the NIH-NCI grants P30CA006973 (L.M. and A.F.), 1U01CA231776 (W.D. and L.M.), and R01CA200859 (W.D. and L.M.), NIH-NIGMS grant R01GM118568 (C.W. and B.L.), R21MH109956-01 (L.C.T. and A.E.J.), and the Department of Defense (DoD) office of the Congressionally Directed Medical Research Programs (CDMRP) award W81XWH-16-1-0739 (E.L.I. and L.M.), Russian Academic project 0112-2019-0001 (A.F.), Fundação de Amparo à Pesquisa do Estado de Minas Gerais award BDS-00493-16 (E.L.I and G.R.F.). *recount2* and `FC-R2` are hosted on SciServer, a collaborative ressearch environment for large-scale data-driven science. It is being developed at, and administered by, the Institute for Data Intensive Engineering and Science (IDIES) at Johns Hopkins University. SciServer is funded by the National Science Foundation Award ACI-1261715. For more information about SciServer, visit http://www.sciserver.org/.

## Author contributions statement

L.M. conceived the idea, L.M., E.L.I., A.F. and B.L. designed the study; E.L.I., D.F.S., T.M., W.D., A.S., L.C.T., and L.M. performed the analysis; E.L.I., D.F.S., F.P.L., G.R.F. and L.M. interpreted the results; L.C.T., C.W., C.Y., K.Y, N.K., M.I., H.S., T.K., C.C.H., M.H., J.W.S., P.C. A.E.J., J.T.L. and B.L. provided the data and tools; E.L.I., D.F.S., L.C.T., B.L. and L.M. wrote the manuscript; All authors reviewed and approved the manuscript.

