## Supplementary Material for "Recounting the FANTOM Cage Associated Transcriptome"

Access to the data and code is available from `http://marchionnilab.org/fcr2.html`, expression data can be directly downloaded from `https://jhubiostatistics.shinyapps.io/recount/` and the *recount* Bioconductor package (v1.9.5 or newer) at `https://bioconductor.org/packages/recount`.

### Supplementary Tables

**Table S1.** Intersection of significant mRNAs across 13 cancer types (Global FDR $< 0.000001$)

| geneID | geneSymbol | geneType | CAT_geneClass | CAT_DHS_type | Chromosome | geneStart | geneEnd | strand | Direction |
|---|---|---|---|---|---|---|---|---|---|
| ENSG00000065328 | MCM10 | protein_coding | coding_mRNA | DHS_promoter | chr10 | 13161579 | 13211916 | + | Up-regulated in tumor |
| ENSG00000090889 | KIF4A | protein_coding | coding_mRNA | DHS_promoter | chrX | 70290040 | 70421061 | + | Up-regulated in tumor |
| ENSG00000091651 | ORC6 | protein_coding | coding_mRNA | DHS_promoter | chr16 | 46689310 | 46702149 | + | Up-regulated in tumor |
| ENSG00000092853 | CLSPN | protein_coding | coding_mRNA | DHS_promoter | chr1 | 35719788 | 35769967 | - | Up-regulated in tumor |
| ENSG00000093009 | CDC45 | protein_coding | coding_mRNA | DHS_promoter | chr22 | 19479466 | 19528387 | + | Up-regulated in tumor |
| ENSG00000100162 | CENPM | protein_coding | coding_mRNA | DHS_promoter | chr22 | 41769677 | 41947123 | - | Up-regulated in tumor |
| ENSG00000102384 | CENPI | protein_coding | coding_mRNA | DHS_promoter | chrX | 101098204 | 101169831 | + | Up-regulated in tumor |
| ENSG00000105011 | ASF1B | protein_coding | coding_mRNA | DHS_promoter | chr19 | 14118278 | 14138467 | - | Up-regulated in tumor |
| ENSG00000106268 | NUDT1 | protein_coding | coding_mRNA | DHS_promoter | chr7 | 2242226 | 2251354 | + | Up-regulated in tumor |
| ENSG00000112984 | KIF20A | protein_coding | coding_mRNA | DHS_promoter | chr5 | 138178724 | 138189480 | + | Up-regulated in tumor |
| ENSG00000115163 | CENPA | protein_coding | coding_mRNA | DHS_promoter | chr2 | 26764321 | 26803376 | + | Up-regulated in tumor |
| ENSG00000117650 | NEK2 | protein_coding | coding_mRNA | DHS_promoter | chr1 | 211658373 | 211675621 | - | Up-regulated in tumor |
| ENSG00000121152 | NCAPH | protein_coding | coding_mRNA | DHS_promoter | chr2 | 96335801 | 96393940 | + | Up-regulated in tumor |
| ENSG00000126787 | DLGAP5 | protein_coding | coding_mRNA | DHS_promoter | chr14 | 55148112 | 55191585 | - | Up-regulated in tumor |
| ENSG00000138180 | CEP55 | protein_coding | coding_mRNA | DHS_promoter | chr10 | 93496639 | 93546919 | + | Up-regulated in tumor |
| ENSG00000144554 | FANCD2 | protein_coding | coding_mRNA | DHS_promoter | chr3 | 10023970 | 10102460 | + | Up-regulated in tumor |
| ENSG00000148773 | MKI67 | protein_coding | coding_mRNA | DHS_promoter | chr10 | 128092566 | 128126423 | - | Up-regulated in tumor |
| ENSG00000156970 | BUB1B | protein_coding | coding_mRNA | DHS_promoter | chr15 | 40161069 | 40223524 | + | Up-regulated in tumor |
| ENSG00000157456 | CCNB2 | protein_coding | coding_mRNA | DHS_promoter | chr15 | 59097077 | 59161020 | + | Up-regulated in tumor |
| ENSG00000162062 | C16orf59 | protein_coding | coding_mRNA | DHS_dyadic | chr16 | 2460109 | 2465114 | + | Up-regulated in tumor |
| ENSG00000165304 | MELK | protein_coding | coding_mRNA | DHS_promoter | chr9 | 36572085 | 36682572 | + | Up-regulated in tumor |
| ENSG00000166508 | MCM7 | protein_coding | coding_mRNA | DHS_promoter | chr7 | 100085474 | 100105533 | - | Up-regulated in tumor |
| ENSG00000167513 | CDT1 | protein_coding | coding_mRNA | DHS_promoter | chr16 | 88799649 | 88818226 | + | Up-regulated in tumor |
| ENSG00000169679 | BUB1 | protein_coding | coding_mRNA | DHS_promoter | chr2 | 110630642 | 110678063 | - | Up-regulated in tumor |
| ENSG00000186185 | KIF18B | protein_coding | coding_mRNA | DHS_promoter | chr17 | 44923253 | 44947773 | - | Up-regulated in tumor |
| ENSG00000189057 | FAM111B | protein_coding | coding_mRNA | DHS_promoter | chr11 | 59063500 | 59131211 | + | Up-regulated in tumor |
| ENSG00000076555 | ACACB | protein_coding | coding_mRNA | DHS_enhancer | chr12 | 109116443 | 109270766 | + | Down-regulated in tumor |
| ENSG00000106034 | CPED1 | protein_coding | coding_mRNA | DHS_promoter | chr7 | 120987841 | 121310413 | + | Down-regulated in tumor |
| ENSG00000112425 | EPM2A | protein_coding | coding_mRNA | DHS_promoter | chr6 | 145382951 | 145763619 | - | Down-regulated in tumor |
| ENSG00000116678 | LEPR | protein_coding | coding_mRNA | DHS_promoter | chr1 | 65420701 | 65643153 | + | Down-regulated in tumor |
| ENSG00000130988 | RGN | protein_coding | coding_mRNA | DHS_promoter | chrX | 47078366 | 47104867 | + | Down-regulated in tumor |
| ENSG00000133800 | LYVE1 | protein_coding | coding_mRNA | DHS_enhancer | chr11 | 10307853 | 10611707 | - | Down-regulated in tumor |
| ENSG00000136842 | TMOD1 | protein_coding | coding_mRNA | DHS_promoter | chr9 | 97501142 | 97628196 | + | Down-regulated in tumor |
| ENSG00000138356 | AOX1 | protein_coding | coding_mRNA | DHS_promoter | chr2 | 200586053 | 200677064 | + | Down-regulated in tumor |

Continued on next page

**Table S1.** Intersection of significant mRNAs across 13 cancer types (Global FDR $< 0.000001$)

| geneID | geneSymbol | geneType | CAT_geneClass | CAT_DHS_type | Chromosome | geneStart | geneEnd | strand | Direction |
|---|---|---|---|---|---|---|---|---|---|
| ENSG00000151623 | NR3C2 | protein_coding | coding_mRNA | DHS_promoter | chr4 | 148077496 | 148445575 | - | Down-regulated in tumor |
| ENSG00000154330 | PGM5 | protein_coding | coding_mRNA | DHS_promoter | chr9 | 68327407 | 68539746 | + | Down-regulated in tumor |
| ENSG00000168546 | GFRA2 | protein_coding | coding_mRNA | DHS_promoter | chr8 | 21365101 | 21812338 | - | Down-regulated in tumor |
| ENSG00000170271 | FAXDC2 | protein_coding | coding_mRNA | DHS_promoter | chr5 | 154810519 | 154859509 | - | Down-regulated in tumor |
| ENSG00000185432 | METTL7A | protein_coding | coding_mRNA | DHS_promoter | chr12 | 50922629 | 50935894 | + | Down-regulated in tumor |
| ENSG00000196616 | ADH1B | protein_coding | coding_mRNA | DHS_enhancer | chr4 | 99286865 | 99321401 | - | Down-regulated in tumor |
| ENSG00000198300 | PEG3 | protein_coding | coding_mRNA | DHS_promoter | chr19 | 56801328 | 56840726 | - | Down-regulated in tumor |

**Table S2.** Intersection of significant divergent promoters across 13 cancer types (Global FDR $< 0.1$)

| geneID | geneSymbol | geneType | CAT_geneClass | CAT_DHS_type | Chromosome | geneStart | geneEnd | strand | Direction |
|---|---|---|---|---|---|---|---|---|---|
| CATG00000017193 | CATG00000017193.1 | __na | lncRNA_divergent | DHS_promoter | chr13 | 52649609 | 52652307 | - | Up-regulated in tumor |
| CATG00000020461 | CATG00000020461.1 | __na | lncRNA_divergent | DHS_promoter | chr14 | 22831988 | 22836822 | - | Up-regulated in tumor |
| CATG00000054098 | CATG00000054098.1 | __na | lncRNA_divergent | DHS_promoter | chr20 | 3043468 | 3048131 | - | Up-regulated in tumor |
| CATG00000087995 | CATG00000087995.1 | __na | lncRNA_divergent | DHS_promoter | chr6 | 30639697 | 30647759 | - | Up-regulated in tumor |
| CATG00000101363 | CATG00000101363.1 | __na | lncRNA_divergent | DHS_promoter | chr8 | 144792564 | 144796174 | + | Up-regulated in tumor |
| ENSG00000228839 | RP3-400N23.6 | antisense | lncRNA_divergent | DHS_promoter | chr22 | 31290906 | 31357952 | + | Up-regulated in tumor |
| ENSG00000235989 | MORC2-AS1 | antisense | lncRNA_divergent | DHS_promoter | chr22 | 30922325 | 30932449 | + | Up-regulated in tumor |
| ENSG00000247373 | RP11-486O12.2 | lincRNA | lncRNA_divergent | DHS_promoter | chr12 | 123575002 | 123584820 | - | Up-regulated in tumor |
| ENSG00000255717 | SNHG1 | processed_transcript | lncRNA_divergent | DHS_promoter | chr11 | 62851874 | 62855885 | - | Up-regulated in tumor |
| ENSG00000257605 | RP11-680A11.5 | antisense | lncRNA_divergent | DHS_promoter | chr12 | 53298394 | 53300360 | - | Up-regulated in tumor |
| ENSG00000258384 | AC068831.6 | antisense | lncRNA_divergent | DHS_promoter | chr15 | 90950782 | 90955229 | - | Up-regulated in tumor |
| ENSG00000260442 | RP11-22P6.3 | antisense | lncRNA_divergent | DHS_promoter | chr16 | 28868384 | 28879920 | - | Up-regulated in tumor |
| ENSG00000263412 | RP5-890E16.2 | processed_transcript | lncRNA_divergent | DHS_promoter | chr17 | 48038796 | 48048670 | - | Up-regulated in tumor |
| ENSG00000270195 | RP11-572O17.1 | lincRNA | lncRNA_divergent | DHS_promoter | chr4 | 1712410 | 1715967 | + | Up-regulated in tumor |
| ENSG00000272455 | RP4-758J18.13 | lincRNA | lncRNA_divergent | DHS_promoter | chr1 | 1407377 | 1410854 | + | Up-regulated in tumor |
| CATG00000105517 | CATG00000105517.1 | __na | lncRNA_divergent | DHS_promoter | chr9 | 70413086 | 70609298 | + | Down-regulated in tumor |
| ENSG00000175611 | LINC00476 | processed_transcript | lncRNA_divergent | DHS_promoter | chr9 | 95759231 | 95875979 | - | Down-regulated in tumor |
| ENSG00000225793 | RP1-234P15.4 | lincRNA | lncRNA_divergent | DHS_promoter | chr6 | 75284598 | 75305766 | + | Down-regulated in tumor |
| ENSG00000226237 | RP11-276H19.1 | lincRNA | lncRNA_divergent | DHS_promoter | chr9 | 86946617 | 87014413 | + | Down-regulated in tumor |
| ENSG00000232160 | RAP2C-AS1 | antisense | lncRNA_divergent | DHS_promoter | chrX | 132217007 | 132435459 | + | Down-regulated in tumor |
| ENSG00000234492 | RPL34-AS1 | lincRNA | lncRNA_divergent | DHS_promoter | chr4 | 108538190 | 108620395 | - | Down-regulated in tumor |
| ENSG00000235652 | RP11-545I5.3 | antisense | lncRNA_divergent | DHS_promoter | chr6 | 145814895 | 145886585 | + | Down-regulated in tumor |

Continued on next page

**Table S2.** Intersection of significant divergent promoters across 13 cancer types (Global FDR $< 0.1$)

| geneID | geneSymbol | geneType | CAT_geneClass | CAT_DHS_type | Chromosome | geneStart | geneEnd | strand | Direction |
|---|---|---|---|---|---|---|---|---|---|
| ENSG00000245293 | RP11-286E11.1 | antisense | lncRNA_divergent | DHS_promoter | chr4 | 107862676 | 107989692 | - | Down-regulated in tumor |
| ENSG00000248866 | USP46-AS1 | lincRNA | lncRNA_divergent | DHS_promoter | chr4 | 52656631 | 52669444 | + | Down-regulated in tumor |
| ENSG00000248980 | RP11-87F15.2 | antisense | lncRNA_divergent | DHS_promoter | chr4 | 176303507 | 176322213 | - | Down-regulated in tumor |
| ENSG00000267414 | RP11-456K23.1 | lincRNA | lncRNA_divergent | DHS_promoter | chr18 | 44664515 | 44680693 | - | Down-regulated in tumor |
| ENSG00000271849 | CTC-332L22.1 | lincRNA | lncRNA_divergent | DHS_promoter | chr5 | 109687802 | 109689251 | - | Down-regulated in tumor |
| ENSG00000272686 | RP11-390E23.6 | antisense | lncRNA_divergent | DHS_promoter | chr7 | 123748542 | 123756392 | + | Down-regulated in tumor |

**Table S3.** Intersection of significant intergenic promoters across 13 cancer types (Global FDR $< 0.1$)

| geneID | geneSymbol | geneType | CAT_geneClass | CAT_DHS_type | Chromosome | geneStart | geneEnd | strand | Direction |
|---|---|---|---|---|---|---|---|---|---|
| ENSG00000196756 | SNHG17 | processed_transcript | lncRNA_intergenic | DHS_promoter | chr20 | 38404766 | 38435328 | - | Up-regulated in tumor |
| ENSG00000260924 | AC004463.6 | antisense | lncRNA_intergenic | DHS_promoter | chr22 | 19171395 | 19175403 | + | Up-regulated in tumor |
| ENSG00000215386 | LINC00478 | lincRNA | lncRNA_intergenic | DHS_promoter | chr21 | 15910380 | 16646424 | + | Down-regulated in tumor |
| ENSG00000263753 | LINC00667 | lincRNA | lncRNA_intergenic | DHS_promoter | chr18 | 5237844 | 5251731 | + | Down-regulated in tumor |

**Table S4.** Intersection of significant enhancers across 13 cancer types (Global FDR $< 0.1$)

| geneID | geneSymbol | geneType | CAT_geneClass | CAT_DHS_type | Chromosome | geneStart | geneEnd | strand | Direction |
|---|---|---|---|---|---|---|---|---|---|
| CATG00000107122 | CATG00000107122.1 | __na | lncRNA_antisense | DHS_enhancer | chr9 | 128118082 | 128124306 | + | Up-regulated in tumor |
| ENSG00000231246 | RP5-965F6.2 | lincRNA | lncRNA_intergenic | DHS_enhancer | chr1 | 112176973 | 112360607 | - | Down-regulated in tumor |
| ENSG00000255958 | RP11-656E20.5 | antisense | lncRNA_antisense | DHS_enhancer | chr12 | 10214161 | 10220555 | - | Down-regulated in tumor |

**Table S5.** Remapping of Andersson's enhancers list to the FANTOM-CAT permissive set. Since originally published, many of enhancers contained in the Andersson's list[35], were reassigned or removed during the assembly of the FANTOM-CAT based on further evidence from the additional transcriptomic datasets used in the meta-assembly, as well as due to the information obtained from orthogonal genomic and epigenomic information, such as DNase I hypersensitivity and other epigenomic marks, as obtained from the Roadmap Epigenomics Project. Based on the new gene models in *FANTOM-CAT*, we verified the overlapping between the original enhancers list and then summarized the results according to current RNA classes in the *FANTOM-CAT*. The counts for the original enhancers that overlap with exons in the FANTOM-CAT gene models are shown in the "Exonic" column on the left. The counts for the enhancers that did not map to any exon, but that were still within the gene boundaries are shown in the "Intronic" column on the right.

| | Exonic | Intronic |
|---|---|---|
| d-lncRNA | 1762 | 1665 |
| i-lncRNA | 1066 | 274 |
| mRNA | 10509 | 7254 |
| other-RNA | 7512 | 2121 |
| pseudogene | 1218 | 208 |
| senseOverlap-RNA | 2631 | 150 |
| small-RNA | 845 | 68 |
| Total | 34282 | 12247 |

**Table S6.** Survival analysis using Cox proportional regression showing the number of e-lncRNAs with prognostic value across the 13 cancer types. *Non-significant* column indicates the number of genes with FDR greater than 0.05. *Cases* represents the number patients at the beginning of follow-up for each tumor type. *Events* is the number of death cases during follow up. *Median time* is given in days.

| Tumor type | Non-significant | FDR $< 0.05$ | Cases | Events | Median time |
|---|---|---|---|---|---|
| Kidney | 13554 | 3850 | 881 | 227 | N.A. |
| Uterus | 16563 | 831 | 596 | 125 | 3365 |
| Stomach | 16850 | 554 | 392 | 158 | 940 |
| Liver | 16970 | 369 | 365 | 130 | 1694 |
| Bladder | 17234 | 153 | 407 | 178 | 1008 |
| Thyroid | 17277 | 111 | 504 | 16 | N.A. |
| Breast | 17305 | 96 | 1080 | 151 | 3941 |
| Colorectal | 17292 | 87 | 602 | 128 | 2532 |
| Lung | 17335 | 69 | 998 | 395 | 1531 |
| Prostate | 17332 | 53 | 496 | 10 | N.A. |
| HeadNeck | 17398 | 3 | 501 | 217 | 1671 |
| Bile | 15725 | 0 | 36 | 18 | 1220 |
| Esophagus | 17404 | 0 | 184 | 77 | 784 |

**Table S7.** Survival analysis using Cox proportional regression showing the number of dp-lncRNAs with prognostic value across the 13 cancer types. *Non-significant* column indicates the number of genes with FDR greater than 0.05. *Cases* represents the number patients at the beginning of follow-up for each tumor type. *Events* is the number of death cases during follow up. *Median time* is given in days.

| Tumor type | Non-significant | FDR $< 0.05$ | Cases | Events | Median time |
|---|---|---|---|---|---|
| Kidney | 4247 | 1957 | 881 | 227 | N.A. |
| Uterus | 5477 | 726 | 596 | 125 | 3365 |
| Liver | 5971 | 223 | 365 | 130 | 1694 |
| Thyroid | 6073 | 128 | 504 | 16 | N.A. |

Continued on next page

**Table S7.** Survival analysis using Cox proportional regression showing the number of dp-lncRNAs with prognostic value across the 13 cancer types. *Non-significant* column indicates the number of genes with FDR greater than 0.05. *Cases* represents the number patients at the beginning of follow-up for each tumor type. *Events* is the number of death cases during follow up. *Median time* is given in days.

| Tumor type | Non-significant | FDR $< 0.05$ | Cases | Events | Median time |
|---|---|---|---|---|---|
| Prostate | 6134 | 69 | 496 | 10 | N.A. |
| Colorectal | 6161 | 41 | 602 | 128 | 2532 |
| Stomach | 6167 | 37 | 392 | 158 | 940 |
| Bladder | 6176 | 27 | 407 | 178 | 1008 |
| Breast | 6195 | 8 | 1080 | 151 | 3941 |
| Lung | 6202 | 2 | 998 | 395 | 1531 |
| Bile | 6078 | 0 | 36 | 18 | 1220 |
| Esophagus | 6204 | 0 | 184 | 77 | 784 |
| HeadNeck | 6204 | 0 | 501 | 217 | 1671 |

**Table S8.** Survival analysis using Cox proportional regression showing the number of ip-lncRNAs with prognostic value across the 13 cancer types. *Non-significant* column indicates the number of genes with FDR greater than 0.05. *Cases* represents the number patients at the beginning of follow-up for each tumor type. *Events* is the number of death cases during follow up. *Median time* is given in days.

| Tumor type | Non-significant | FDR $< 0.05$ | Cases | Events | Median time |
|---|---|---|---|---|---|
| Kidney | 1471 | 477 | 881 | 227 | N.A. |
| Uterus | 1660 | 287 | 596 | 125 | 3365 |
| Liver | 1864 | 78 | 365 | 130 | 1694 |
| Colorectal | 1896 | 46 | 602 | 128 | 2532 |
| Thyroid | 1909 | 37 | 504 | 16 | N.A. |
| Stomach | 1919 | 29 | 392 | 158 | 940 |
| Bladder | 1930 | 18 | 407 | 178 | 1008 |
| Prostate | 1929 | 18 | 496 | 10 | N.A. |
| Breast | 1931 | 17 | 1080 | 151 | 3941 |
| Lung | 1938 | 9 | 998 | 395 | 1531 |
| HeadNeck | 1944 | 3 | 501 | 217 | 1671 |
| Bile | 1866 | 0 | 36 | 18 | 1220 |
| Esophagus | 1947 | 0 | 184 | 77 | 784 |

**Table S9.** Survival analysis using Cox proportional regression showing the number of mRNAs with prognostic value across the 13 cancer types. *Non-significant* column indicates the number of genes with FDR greater than 0.05. *Cases* represents the number patients at the beginning of follow-up for each tumor type. *Events* is the number of death cases during follow up. *Median time* is given in days.

| Tumor type | Non-significant | FDR $< 0.05$ | Cases | Events | Median time |
|---|---|---|---|---|---|
| Kidney | 12933 | 9166 | 881 | 227 | N.A. |
| Uterus | 16698 | 5400 | 596 | 125 | 3365 |
| Liver | 18521 | 3571 | 365 | 130 | 1694 |
| Prostate | 21458 | 638 | 496 | 10 | N.A. |
| Colorectal | 21641 | 455 | 602 | 128 | 2532 |

Continued on next page

**Table S9.** Survival analysis using Cox proportional regression showing the number of mRNAs with prognostic value across the 13 cancer types. *Non-significant* column indicates the number of genes with FDR greater than 0.05. *Cases* represents the number patients at the beginning of follow-up for each tumor type. *Events* is the number of death cases during follow up. *Median time* is given in days.

| Tumor type | Non-significant | FDR $< 0.05$ | Cases | Events | Median time |
|---|---|---|---|---|---|
| Bladder | 21674 | 424 | 407 | 178 | 1008 |
| Thyroid | 21721 | 374 | 504 | 16 | N.A. |
| Stomach | 21797 | 302 | 392 | 158 | 940 |
| Lung | 21922 | 177 | 998 | 395 | 1531 |
| Breast | 21998 | 101 | 1080 | 151 | 3941 |
| HeadNeck | 22020 | 78 | 501 | 217 | 1671 |
| Bile | 21952 | 0 | 36 | 18 | 1220 |
| Esophagus | 22099 | 0 | 184 | 77 | 784 |

**Table S10.** Differentially expressed mRNA genes with prognostic value across cancer types. *Good* indicates the Cox HR $< 1$. *Bad* represents Cox HR $> 1$. *N.P.* refers that the given gene is non-prognostic on this cancer type

| | Bladder | Breast | Colorectal | HeadNeck | Kidney | Liver | Lung | Prostate | Stomach | Thyroid | Uterus |
|---|---|---|---|---|---|---|---|---|---|---|---|
| ENSG00000065328 | N.P. | N.P. | N.P. | N.P. | Bad | Bad | N.P. | Bad | N.P. | N.P. | Bad |
| ENSG00000090889 | N.P. | N.P. | N.P. | N.P. | Bad | Bad | N.P. | Bad | N.P. | N.P. | Bad |
| ENSG00000091651 | N.P. | N.P. | N.P. | N.P. | Bad | Bad | N.P. | Bad | N.P. | N.P. | N.P. |
| ENSG00000092853 | N.P. | N.P. | N.P. | N.P. | Bad | Bad | N.P. | Bad | N.P. | N.P. | N.P. |
| ENSG00000093009 | N.P. | N.P. | N.P. | N.P. | Bad | Bad | N.P. | N.P. | N.P. | N.P. | N.P. |
| ENSG00000100162 | N.P. | N.P. | N.P. | N.P. | Bad | Bad | N.P. | N.P. | N.P. | N.P. | N.P. |
| ENSG00000102384 | N.P. | N.P. | N.P. | N.P. | Bad | Bad | N.P. | Bad | N.P. | N.P. | Bad |
| ENSG00000105011 | N.P. | N.P. | N.P. | N.P. | Bad | Bad | N.P. | N.P. | N.P. | N.P. | N.P. |
| ENSG00000106268 | N.P. | N.P. | N.P. | N.P. | Bad | Bad | N.P. | N.P. | N.P. | N.P. | N.P. |
| ENSG00000112984 | N.P. | N.P. | N.P. | N.P. | Bad | Bad | N.P. | N.P. | N.P. | N.P. | Bad |
| ENSG00000115163 | N.P. | N.P. | N.P. | N.P. | Bad | Bad | N.P. | N.P. | N.P. | N.P. | Bad |
| ENSG00000117650 | N.P. | N.P. | N.P. | N.P. | Bad | Bad | N.P. | N.P. | N.P. | N.P. | N.P. |
| ENSG00000121152 | N.P. | N.P. | N.P. | N.P. | Bad | Bad | N.P. | N.P. | N.P. | N.P. | Bad |
| ENSG00000126787 | N.P. | N.P. | N.P. | N.P. | Bad | Bad | N.P. | N.P. | N.P. | N.P. | Bad |
| ENSG00000138180 | N.P. | N.P. | N.P. | N.P. | Bad | Bad | N.P. | N.P. | N.P. | N.P. | N.P. |
| ENSG00000144554 | N.P. | N.P. | N.P. | N.P. | Bad | Bad | N.P. | N.P. | N.P. | N.P. | N.P. |
| ENSG00000148773 | N.P. | N.P. | N.P. | N.P. | Bad | Bad | N.P. | N.P. | N.P. | N.P. | N.P. |
| ENSG00000156970 | N.P. | N.P. | N.P. | N.P. | Bad | Bad | N.P. | Bad | N.P. | N.P. | Bad |
| ENSG00000157456 | N.P. | N.P. | N.P. | N.P. | Bad | Bad | N.P. | Bad | N.P. | N.P. | N.P. |
| ENSG00000162062 | N.P. | N.P. | N.P. | N.P. | Bad | Bad | N.P. | N.P. | N.P. | N.P. | Bad |
| ENSG00000165304 | N.P. | N.P. | N.P. | N.P. | Bad | Bad | N.P. | N.P. | N.P. | N.P. | N.P. |
| ENSG00000166508 | N.P. | N.P. | N.P. | N.P. | Bad | N.P. | N.P. | N.P. | N.P. | N.P. | Bad |
| ENSG00000167513 | N.P. | N.P. | N.P. | N.P. | Bad | Bad | N.P. | N.P. | N.P. | N.P. | N.P. |
| ENSG00000169679 | N.P. | N.P. | N.P. | N.P. | Bad | Bad | N.P. | N.P. | N.P. | N.P. | Bad |
| ENSG00000186185 | N.P. | N.P. | N.P. | N.P. | Bad | Bad | N.P. | Bad | N.P. | N.P. | Bad |
| ENSG00000189057 | N.P. | N.P. | N.P. | N.P. | Bad | Bad | N.P. | N.P. | N.P. | N.P. | N.P. |
| ENSG00000076555 | N.P. | N.P. | N.P. | N.P. | Good | N.P. | N.P. | N.P. | N.P. | N.P. | Bad |
| ENSG00000112425 | N.P. | N.P. | N.P. | N.P. | Good | N.P. | N.P. | N.P. | N.P. | N.P. | Bad |
| ENSG00000130988 | N.P. | N.P. | N.P. | N.P. | Good | Good | N.P. | N.P. | N.P. | N.P. | Bad |
| ENSG00000133800 | N.P. | N.P. | N.P. | N.P. | N.P. | N.P. | N.P. | N.P. | N.P. | N.P. | Bad |

Continued on next page

**Table S10.** Differentially expressed mRNA genes with prognostic value across cancer types. *Good* indicates the Cox HR $< 1$. *Bad* represents Cox HR $> 1$. *N.P.* refers that the given gene is non-prognostic on this cancer type

| | Bladder | Breast | Colorectal | HeadNeck | Kidney | Liver | Lung | Prostate | Stomach | Thyroid | Uterus |
|---|---|---|---|---|---|---|---|---|---|---|---|
| ENSG00000138356 | N.P. | N.P. | N.P. | N.P. | N.P. | N.P. | N.P. | N.P. | N.P. | N.P. | Bad |
| ENSG00000151623 | N.P. | N.P. | N.P. | N.P. | Good | N.P. | N.P. | N.P. | N.P. | N.P. | N.P. |
| ENSG00000154330 | N.P. | N.P. | N.P. | N.P. | Good | N.P. | N.P. | N.P. | N.P. | N.P. | N.P. |
| ENSG00000168546 | N.P. | N.P. | N.P. | N.P. | Bad | N.P. | N.P. | N.P. | N.P. | N.P. | Bad |
| ENSG00000170271 | N.P. | N.P. | N.P. | N.P. | Good | N.P. | N.P. | N.P. | N.P. | Bad | N.P. |
| ENSG00000185432 | N.P. | N.P. | N.P. | N.P. | Good | N.P. | N.P. | N.P. | N.P. | N.P. | N.P. |
| ENSG00000198300 | N.P. | N.P. | N.P. | N.P. | N.P. | N.P. | N.P. | N.P. | N.P. | Bad | N.P. |

**Table S11.** Differentially expressed e-lncRNA genes with prognostic value across cancer types. *Good* indicates the Cox HR $< 1$. *Bad* represents Cox HR $> 1$. *N.P.* refers that the given gene is non-prognostic on this cancer type

| | Bladder | Breast | Colorectal | HeadNeck | Kidney | Liver | Lung | Prostate | Stomach | Thyroid | Uterus |
|---|---|---|---|---|---|---|---|---|---|---|---|
| CATG00000107122 | N.P. | N.P. | N.P. | N.P. | Bad | N.P. | N.P. | N.P. | N.P. | N.P. | N.P. |
| ENSG00000255958 | N.P. | N.P. | N.P. | N.P. | N.P. | N.P. | N.P. | N.P. | Bad | N.P. | N.P. |

**Table S12.** Differentially expressed dp-lncRNA genes with prognostic value across cancer types. *Good* indicates the Cox HR $< 1$. *Bad* represents Cox HR $> 1$. *N.P.* refers that the given gene is non-prognostic on this cancer type

| | Bladder | Breast | Colorectal | Kidney | Liver | Lung | Prostate | Stomach | Thyroid | Uterus |
|---|---|---|---|---|---|---|---|---|---|---|
| CATG00000017193 | N.P. | N.P. | N.P. | Bad | N.P. | N.P. | N.P. | N.P. | N.P. | N.P. |
| CATG00000020461 | N.P. | N.P. | N.P. | Bad | N.P. | N.P. | N.P. | N.P. | N.P. | N.P. |
| CATG00000054098 | N.P. | N.P. | N.P. | Bad | N.P. | N.P. | N.P. | N.P. | N.P. | N.P. |
| CATG00000087995 | N.P. | N.P. | N.P. | N.P. | N.P. | N.P. | Bad | N.P. | N.P. | N.P. |
| CATG00000101363 | N.P. | N.P. | N.P. | Bad | N.P. | N.P. | N.P. | N.P. | N.P. | Bad |
| ENSG00000235989 | N.P. | N.P. | Bad | Bad | N.P. | N.P. | N.P. | N.P. | N.P. | Bad |
| ENSG00000247373 | N.P. | N.P. | N.P. | Bad | N.P. | N.P. | N.P. | N.P. | N.P. | N.P. |
| ENSG00000255717 | N.P. | N.P. | N.P. | Bad | Bad | N.P. | N.P. | N.P. | N.P. | Bad |
| ENSG00000257605 | N.P. | N.P. | N.P. | Bad | N.P. | N.P. | N.P. | N.P. | N.P. | Bad |
| ENSG00000258384 | N.P. | N.P. | N.P. | Bad | N.P. | N.P. | N.P. | N.P. | N.P. | N.P. |
| ENSG00000260442 | N.P. | N.P. | N.P. | Bad | N.P. | N.P. | N.P. | N.P. | N.P. | N.P. |
| ENSG00000263412 | N.P. | N.P. | N.P. | Bad | N.P. | N.P. | N.P. | N.P. | N.P. | Bad |
| ENSG00000270195 | N.P. | N.P. | N.P. | Bad | N.P. | N.P. | N.P. | N.P. | N.P. | N.P. |
| ENSG00000272455 | N.P. | N.P. | N.P. | Bad | N.P. | N.P. | N.P. | N.P. | N.P. | Bad |
| CATG00000105517 | N.P. | N.P. | N.P. | N.P. | N.P. | N.P. | N.P. | N.P. | N.P. | Bad |
| ENSG00000226237 | N.P. | N.P. | N.P. | Bad | N.P. | N.P. | N.P. | N.P. | N.P. | Bad |
| ENSG00000232160 | N.P. | N.P. | N.P. | Good | N.P. | N.P. | N.P. | N.P. | N.P. | N.P. |
| ENSG00000234492 | N.P. | N.P. | N.P. | Good | N.P. | N.P. | N.P. | N.P. | N.P. | N.P. |
| ENSG00000235652 | N.P. | N.P. | N.P. | Good | N.P. | N.P. | N.P. | N.P. | N.P. | N.P. |
| ENSG00000248980 | N.P. | N.P. | N.P. | Good | N.P. | N.P. | N.P. | N.P. | N.P. | N.P. |
| ENSG00000267414 | N.P. | N.P. | N.P. | Good | N.P. | N.P. | N.P. | N.P. | N.P. | Bad |
| ENSG00000271849 | N.P. | N.P. | N.P. | Good | N.P. | N.P. | N.P. | N.P. | N.P. | N.P. |

**Table S13.** Differentially expressed ip-lncRNA genes with prognostic value across cancer types. *Good* indicates the Cox HR $< 1$. *Bad* represents Cox HR $> 1$. *N.P.* refers that the given gene is non-prognostic on this cancer type

| | Bladder | Breast | Colorectal | HeadNeck | Kidney | Liver | Lung | Prostate | Stomach | Thyroid | Uterus |
|---|---|---|---|---|---|---|---|---|---|---|---|
| ENSG00000196756 | N.P. | N.P. | N.P. | N.P. | Bad | N.P. | N.P. | Bad | N.P. | N.P. | Bad |
| ENSG00000260924 | N.P. | N.P. | N.P. | N.P. | Bad | N.P. | N.P. | N.P. | N.P. | N.P. | N.P. |
| ENSG00000215386 | N.P. | N.P. | N.P. | N.P. | N.P. | N.P. | N.P. | N.P. | N.P. | N.P. | Bad |
| ENSG00000263753 | N.P. | N.P. | N.P. | N.P. | N.P. | N.P. | N.P. | N.P. | N.P. | Bad | Bad |

#### Supplementary Figures

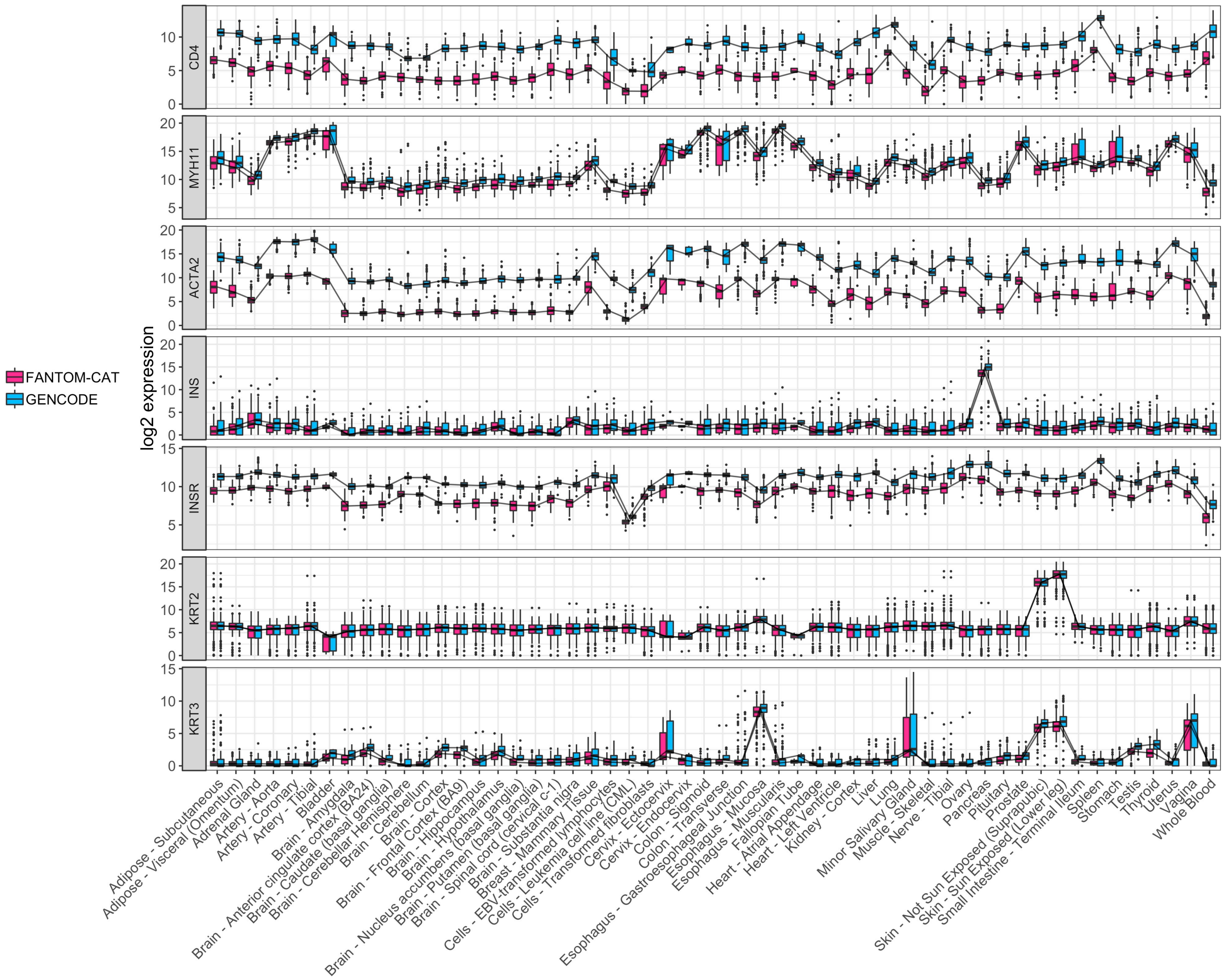

**Figure S1. Tissue specific expression in GTEx.** Log2 expression for tissue specific genes in GTEx data stratify by tissue type using `FC-R2` and GENCODE based quantification. Expression profiles are highly correlated and expressed consistently in the expected tissue types. *CD4* is most expressed in spleen, *MYH11* and *ACTA2* in tissues consisted of muscle cells, *INS* in pancreas and its receptor *INSR* globally, *KRT2* and *KRT3* in skin.

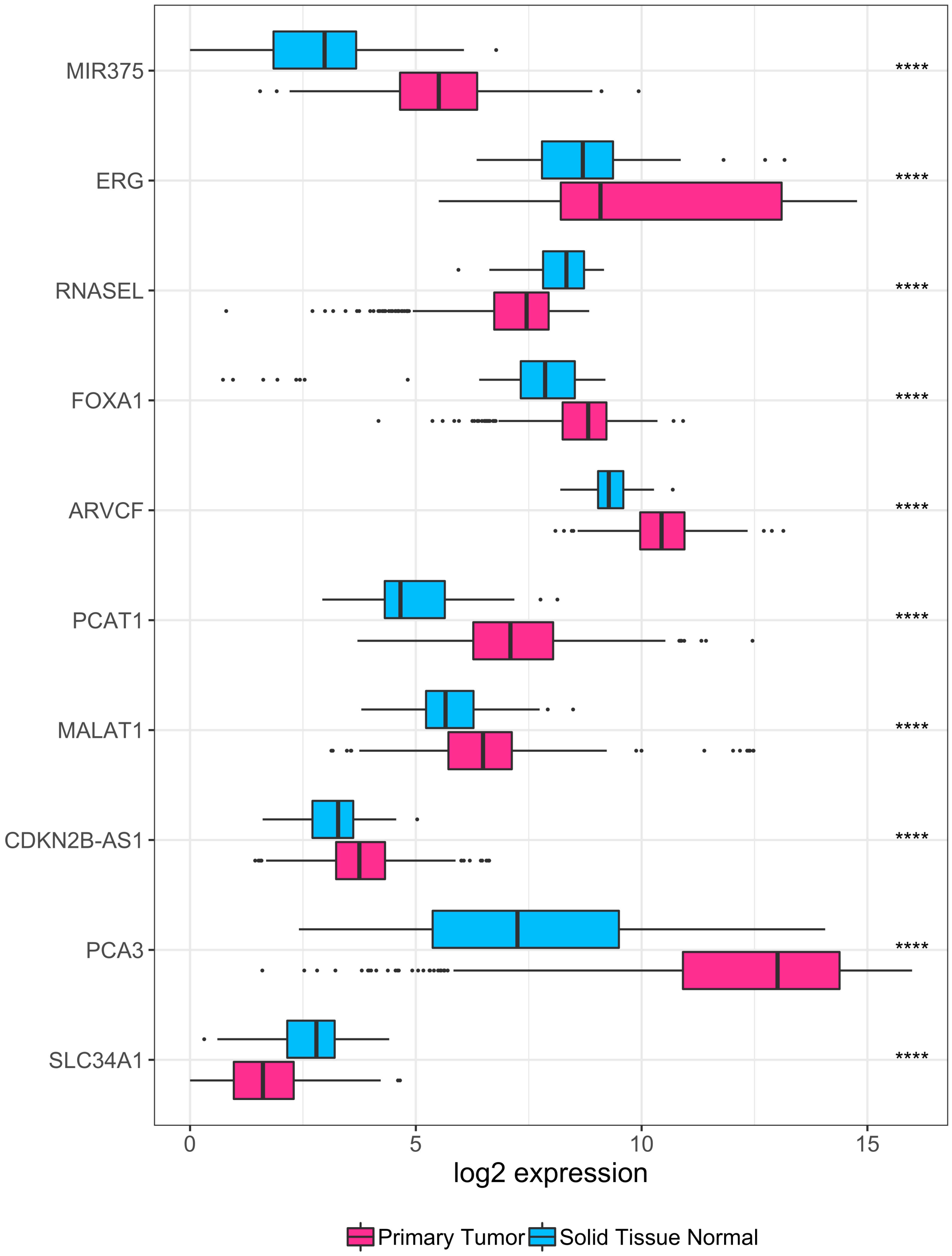

**Figure S2. Coding and non-coding RNAs in prostate cancer.** Expression levels for coding and non-coding RNAs known to be involved in prostate cancer. Results are stratified by sample type, pink is used for normal primary tissue, while light blue for primary cancers. The stars indicate significance from the moderated t-test ($FDR < 0.0001$).

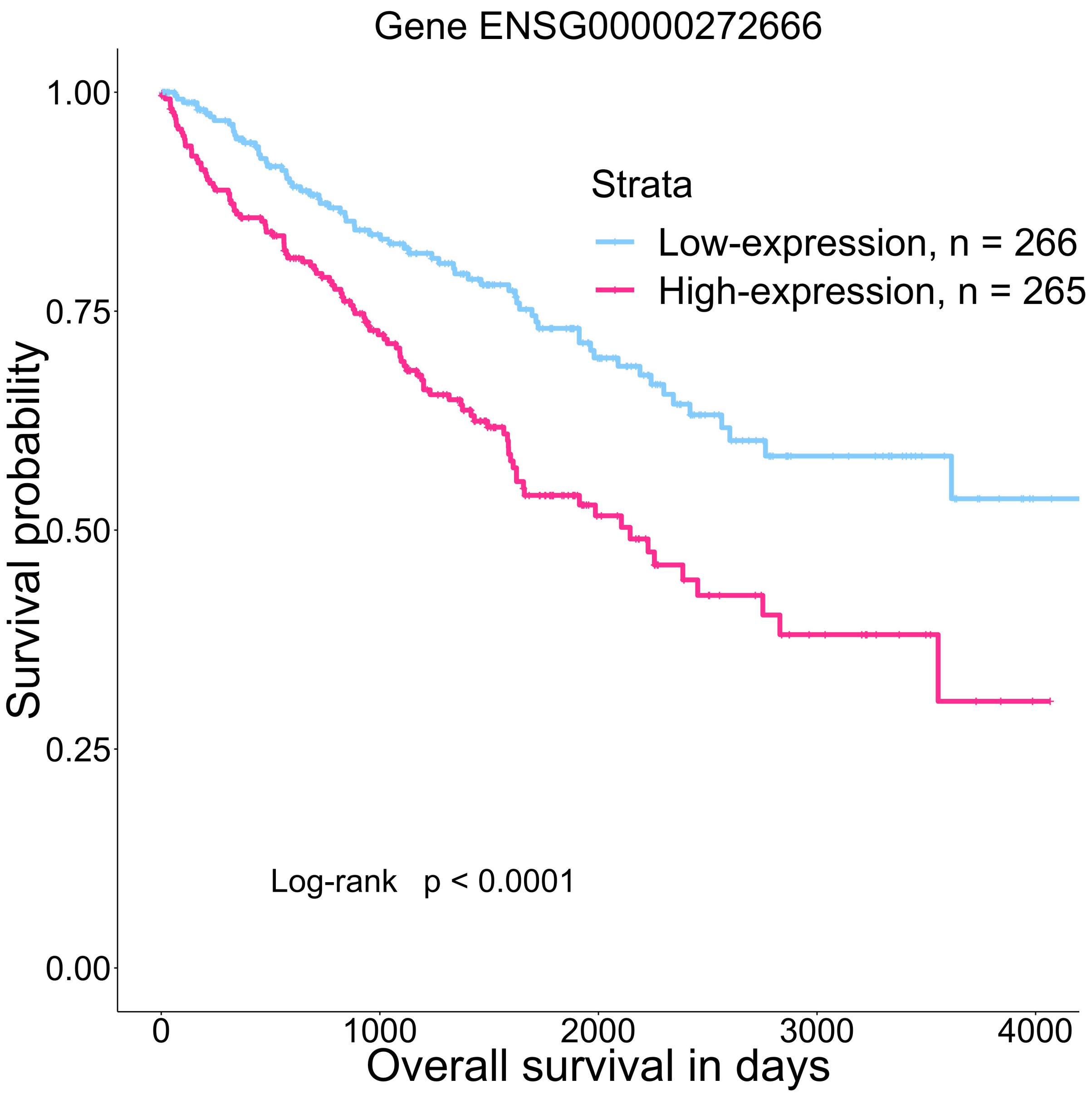

**Figure S3. KM curve for enhancer 22.** Kaplan-Meier survival curve depicting ENSG00000272666 (enhancer22 in Chen et al.[34]) groups split by median expression level for kidney clear cell renal cell carcinoma (KIRC) cases. Statistical significance were assessed using log-rank test.

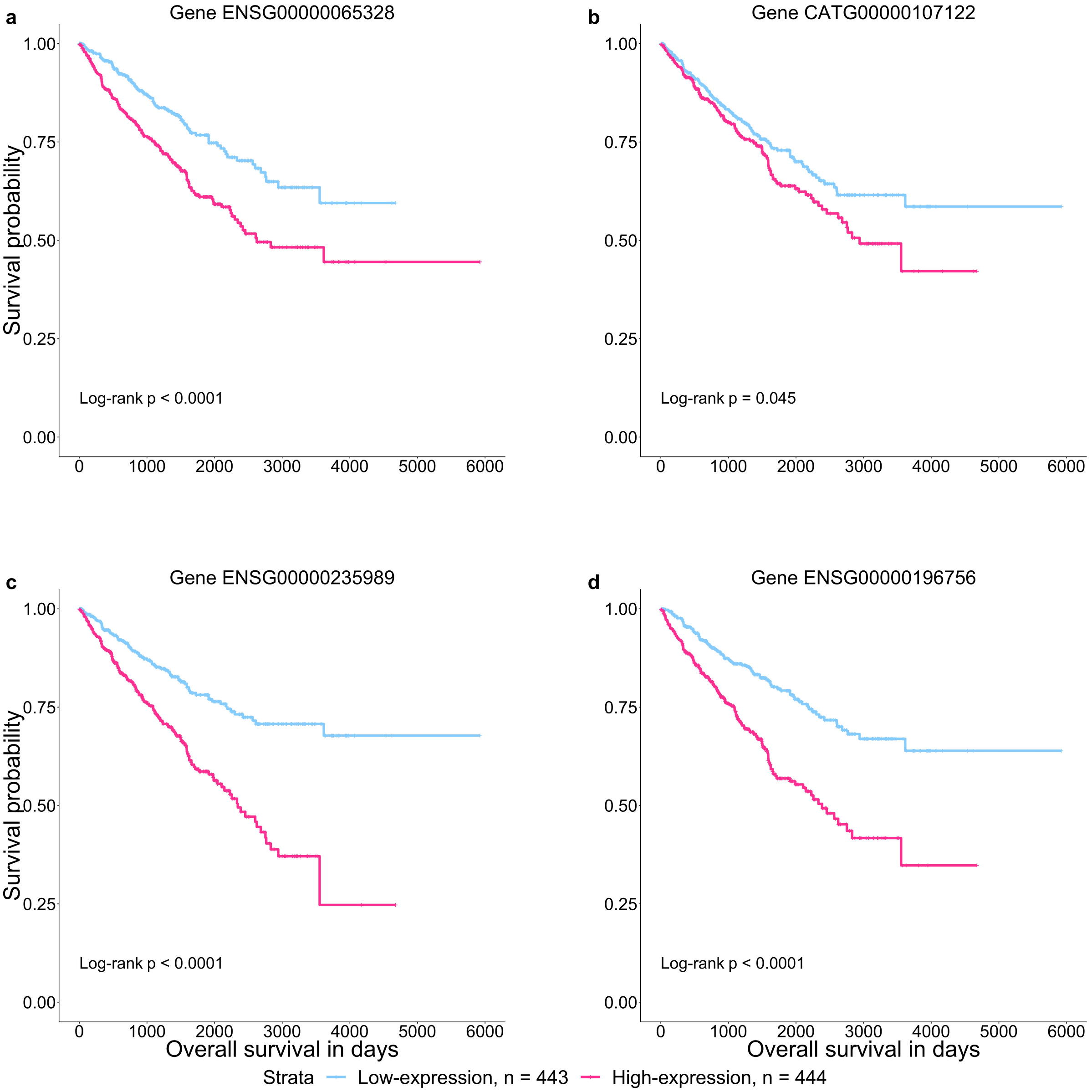

**Figure S4. Kaplan-Meier survival curves depicting four selected differentially expressed genes holding prognostic value in kidney cancers. a.** mRNA gene ENSG00000065328 (*MCM10*), **b.** e-lncRNA gene CATG00000107122 unique to FANTOM-CAT, **c.** dp-lncRNA gene ENSG00000235989 (*MORC2-AS1*), **d.** ip-lncRNA gene ENSG00000196756 (*SNHG17*). Low- and high-expression groups were split by median expression level for each gene. Statistical significance were assessed using log-rank test.

#### Supplementary Methods

##### Data and pre-processing.

We obtained files containing coordinates of updated gene models of FANTOM-CAT permissive set from the ongoing FANTOM6 consortium. These files containing coordinates for 709,176 transcript models were imported into an R session and used to create a Genomic Range object with the GenomicRanges package[44] of all exons coordinates. Given the unstranded nature of recount2 we opted to remove overlapping exons belonging to different gene models to avoid over-quantification of these regions. To avoid losing strand information from annotation we first split exons coordinates by strand into two objects. We then used the `disjoin` function from Genomic Ranges package to generate disjoint segments in each strand, and then across strands. (Figure 5). Each segment coordinate was then assigned to the corresponding overlapping gene model. Segments that were assigned to more than one gene were discarded. The expression values for each segment was then quantified using *recount.bwtool*[45] (code at `https://github.com/LieberInstitute/marchionni_projects`). The resulting expression quantifications were processed to generate `RangedSummarizedExperiment` objects compatible with the *recount2* framework[7,43] (code at `https://github.com/eddieimada/fcr2`). The expression values for each segment was added to its respective gene model, resulting in the final object containing expression at gene level, which is distributed through the recount package and recount2 website.

##### Correlation with other studies.

Due to the decision of removing segments overlapping with more than one gene model, we further investigated if removal of these segments caused significant impact on expression levels. To achieve that we compared our GTEx counts tables to the published GTEx counts from *recount2*. The version 2 of the gene counts for the GTEx samples were downloaded from the recount website (`https://jhubiostatistics.shinyapps.io/recount/`). Next, we scaled each object to a 40M depth using the `scale_counts` function from recount package. After scalling, we obtained the intersection of the genes across both objects and computed the Pearson correlation for each gene. We further selected a few tissue markers to evaluate expression specificity across tissue types.

##### Expression specificity of tissue facets.

To analyze the expression level and specificity of each gene, we first scaled GTEx data to 40M depth using the `scale_counts` function from recount package. Genes were then stratified by RNA class (i.e. mRNA, e-lncRNA, dp-lncRNA, ip-lncRNA) and grouped by tissue type (n = 54 facets). The expression level for each gene was represented by the maximum transcripts per million (TPM) of all samples within a facet. The expression specificity was calculated as the empirical entropy of the mean expression values of each facet divide by the log2 of the number of facets, as follows

$$SPECIFICITY = 1 - (entropy(X)/log_2(N))$$

Where $X$ is a vector of sample-average values for a given gene over all facets and $N$ is the number of facets. The 99.99 percent confidence intervals for the expression of each category by facet were calculated based on TPM values. Genes with a TPM greater than 0.01 were considered expressed.

##### Identification of differentially expressed genes.

To perform differential gene expression analysis across cancer types we relied on TCGA data scaled to 40M depth using the `scale_counts` function from recount. We split each cancer dataset by RNA class (coding mRNA, intergenic promoter lncRNA, divergent promoter lncRNA and enhancer lncRNA) and removed all metastatic samples prior analysis. Each RNA class was treated independently.

The design matrices were created from a factor with two levels (Primary Tumor and Normal Tissue) by setting the normal tissues as the intercept. For each RNA class we removed genes with low expression ( $< 5$ counts) in more than 1/3 of the total samples in each cancer type. After filtering, we normalized raw libraries sizes using method TMM with `calcNormFactors` from edgeR package. After normalization, we transformed the count data to log2-counts per million (logCPM), and estimated the mean-variance relationship to compute appropriate observation-level weights using `voom`.

Finally, we run surrogate variable analysis (SVA)[47] using the permutation procedure proposed by Buja and Eyuboglu[49] and added the first three most variable coefficients as covariates in our design matrix. A generalized linear model approach coupled with empirical Bayes standard errors[46] was then used for identifying differentially expressed genes between the samples. Correction for multiple testing was performed across RNA classes by merging the resulting p-values for each cancer type and applying the Benjamini-Hochberg method[48].

##### Prognostic enhancers analysis.

Univariate Cox proportional regression were performed in four RNA subtypes (22106 mRNAs, 17,404 e-lncRNAs, 6,204 dp-lncRNAs, and 1,948 ip-lncRNAs) available through `FC-R2` as predictors on each of the 13 TCGA cancer types with available survival follow-up. The gene expressions were obtained across cancer types on TCGA data scaled to 40M depth using the `scale_counts` function from recount. We split each cancer dataset by RNA class (coding mRNA, intergenic promoter lncRNA, divergent promoter lncRNA and enhancer lncRNA) using only primary tumor samples gene expression. Some patients had more than one sample, in such cases, the first sample was chosen for the gene expression levels. Each RNA class was treated independently. Survival analysis including Cox proportional regression was done using R survival package. Survival data (follow-up and vital status) available in the phenotype data derived from *recount2 expressionSets* were used to create survival objects for each of the 13 cancer types. Cox proportional regression was done for each cancer type using one gene at a time (univariate regression) grouped on each of the four RNA categories surveyed. Genes were considered predictive if its FDR were equal or less than 0.05, using Benjamini-Hochberg[48] correction. The correction was done within a cancer type grouping all genes by RNA type (i.e. corrected for 22,106 genes if mRNA were surveyed). This procedure yielded the predictive genes list broken-down in the four RNA categories reported. For reporting differentially expressed genes (DEG) among all cancer types that portrait predictive value, the DEG lists were surveyed on each of the significant prognostic genes lists generated during the univariate Cox analysis by cancer types and summary tables reporting the genes predictive potential as good (HR $< 1$), bad (HR $> 1$), and non-predictive (FDR $> 0.05$) were prepared. Kaplan-Meier curves were done using *survminer* R package[50]. Groups were defined using median gene expression level and significance were assessed by log-rank test (P-value $<$ 0.05).

Supplementary data from Chen et al.[34] containing enhancers position and prognostic potential were obtained from the original publication supplementary material. Liftover to hg38 genome assembly was performed to match `FC-R2` coordinates in order to compare both resources. Prognostic genes list provided in Chen's paper were compared to prognostic genes obtained using `FC-R2`.
